# On multistability and constitutive relations of motion of MDA-MB-231 cells on Fibronectin lanes

**DOI:** 10.1101/2022.08.30.505377

**Authors:** Behnam Amiri, Johannes C.J. Heyn, Christoph Schreiber, Joachim O. Rädler, Martin Falcke

## Abstract

Cell motility on flat substrates exhibits multistability between steady and oscillatory morphodynamics, spread and moving states, the biphasic adhesion-velocity relation, and the universal correlation between speed and persistence (UCSP) as simultaneously observed phenomena. Their universality and concurrency suggest a unifying mechanism to exist causing all of them. We search for that mechanism by investigating trajectories of MDA-MB-231 cells on Fibronectin lanes. We also find multistability caused by the clutch mechanism of retrograde flow. Control of the clutch parameters by integrin signalling causes the biphasic adhesion-velocity relation. Protrusion competition based on the clutch causes direction reversal events, the statistics of which explains the UCSP. We suggest that F-actin polymerisation, clutch mechanism of retrograde flow, protrusion competition via membrane tension and drag forces cause the multistability and dynamic cell states, state transition statistics causes the UCSP and the control of this dynamic system by integrin signalling entails the adhesion-velocity relation.

## 1 Introduction

The motion of eukaryotic cells is essential for embryonic development, wound healing, immune responses, and tumour metastasis [1]. Much effort has been devoted to the study of mesenchymal migration with prototypical *in vitro* motion of cells on 2-dimensional adhesive substrates. Cell migration starts with polarisation breaking the spatial symmetry and the formation of a lamellipodium, which is a protrusion of a thin sheet of cytoplasm (0.1–0.3 μm thick) covering tens to hundreds of square micrometres [2–7]. The lamellipodium is mechanically stabilised by adhesion with the substrate [8–13] and is constructed from a network of actin filaments [14–18]. Polymerisation of filament barbed ends at the leading edge of the lamellipodium generates motion and pushes the edge forward [19–22]. Further back, the pointed ends depolymerise and replenish the pool of actin monomers [17, 18]. Once cells are moving, their shape is determined by internal force generation patterns and adhesion [23–28].

Many cell types obey both the adhesion-velocity relation and the universal correlation between speed and persistence (UCSP). The dependency of the cell velocity on adhesion exhibits a velocity maximum at intermediate strength, and slower velocities both at weak and strong adhesion [9–13, 25, 29–33]. Results on the UCSP, describing the relation between cell velocity and persistence time, suggest it to be of similar universality [34]. The faster cells move the more persistent they move. Maiuri et al. report this observation for many different cell types and suggest persistence time to depend exponentially on cell velocity [34]. These types of relations describing the response of a system to external parameters are called constitutive relations in Physics and Engineering. The stationary force-velocity relation is another constitutive relation we will discuss.

Another general observation is that both the shape and the motile state of cells is highly dynamic. Cells stop and start to move again, develop new protrusions, and change direction [24, 28, 35–50]. In addition to these states of motion, there exist states distinguished by the dynamic regime of front protrusion and cell back and/or back protrusion. Stationary and oscillatory dynamic regimes with one or several protrusions have been observed, and caused a surge of interest in multistability in cell motility [5, 38, 42, 43, 51–59].

Multistability with its state dynamics, biphasic adhesion-velocity relation and the UCSP appear to describe the motile behaviour of many different cell types. Mechanisms have been suggested for multistability [38, 51, 52, 55, 56, 58, 59], the biphasic adhesion-velocity relation [10, 29, 30, 33], the UCSP [34] and the stationary force-velocity relation [60] each separately. However, the generality and concurrency of multistability and the constitutive relations strongly suggest that a single mechanism should explain all three of them.

In search for such a mechanism, we carry out a series of experiments with MDA-MB-231 cells on 1d lanes in a range of Fibronectin concentrations and formulate our suggestion for a mechanism as a biophysical model based on the studies [30, 61]. The mechanism involves the force balance at the protrusion edges, the clutch mechanism of retrograde flow, competing protrusions and integrin signalling. Our key finding is that the intracellular dynamics generating multistability also determine the constitutive relations. We will introduce basic experimental observations in section 2.1, characterise cell states and compare experimental and simulated data in section 2.2 and explain state transitions and their relation to the UCSP in section 2.3. We will explain the ideas defining the theory and compute the force-velocity relation and adhesion-velocity relation in section 4.6.

## 2 Results

### 2.1 Dynamic cell states

We monitored MDA-MB-231 cells migrating in Fibronectin coated lanes using scanning time-lapse microscopy. Movies from several fields of view were recorded with typically 10 min intervals over 48 h collecting in total more than 20.000 cell trajectories as described in the methods section. A representative kymograph of the 1-dimensional cell motion is depicted in Fig. 1A. We find 4 different dynamic states (Fig. 1B-E): a spread state with steady length (SS), a spread state with oscillatory protrusions at both ends (SO), a moving state with steady length (MS), and a moving state with an oscillating back protrusion (MO). The two moving states exist as up and down going, so that we observe 6 states in total. We also exemplify 6 different state transitions. All of them occur on homogeneous Fibronectin lanes and without any stimulation. Therefore, we call them spontaneous transitions. The states SS and MO have also been found with RPE1 cells and NIH-3T3 fibroblasts [42], and all four dynamic states with C6 glioma cells.

**Figure 1:**
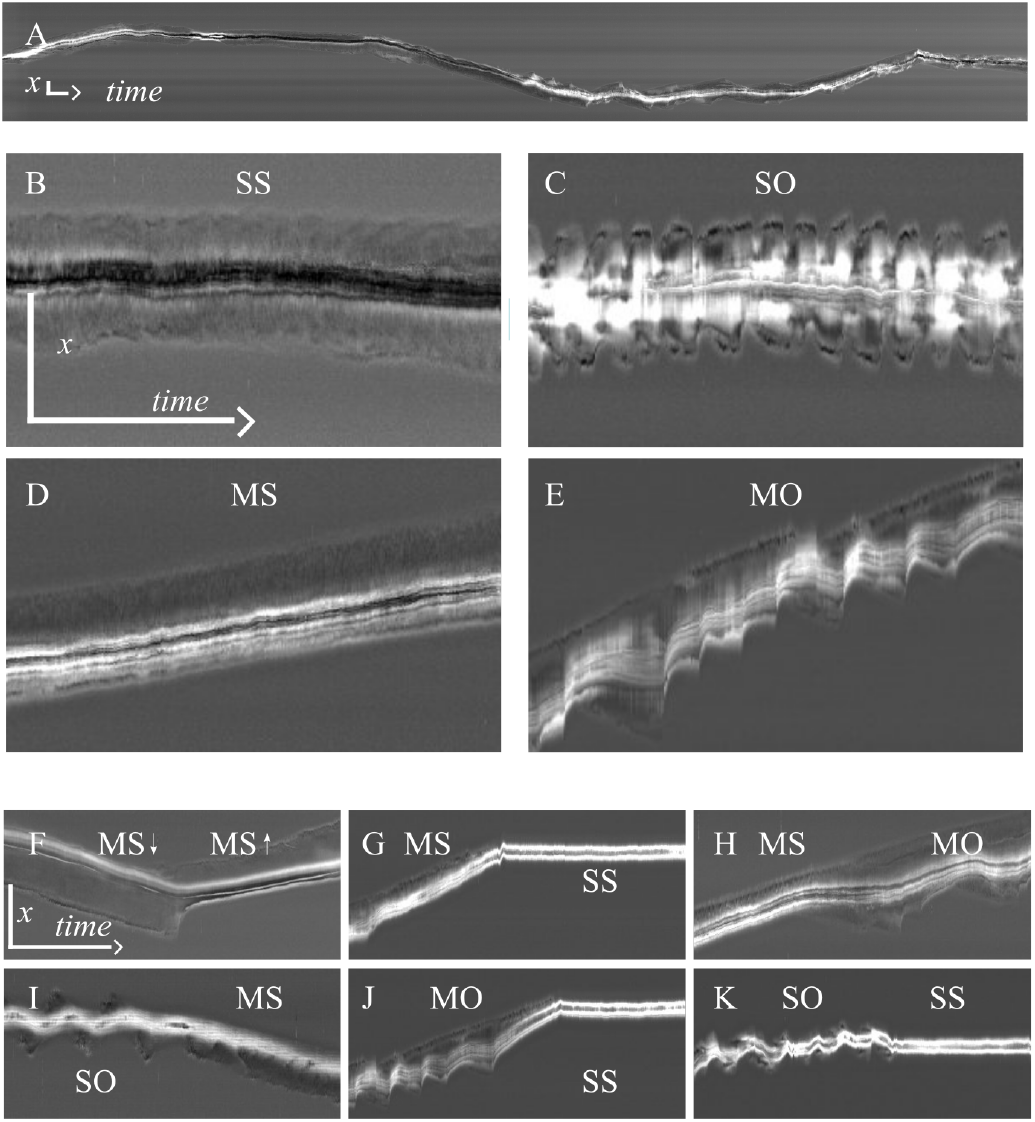
Four dynamic states and two motion directions form the multistability of MDA-MB-231 cells on homogeneous Fibronectin lanes. (A) Kymograph from a typical 48h-trajectory. (B) A spread (S) cell with steady length (S). We call this state SS. (C) A spread cell with oscillatory length (O). We call this state SO. (D) A moving cell (M) with steady length - state MS. (E) A moving cell with oscillating length - state MO. (F) Transition from a downward moving MS state to an upward moving MS state. (G)-(K) The cells undergo a MS→SS, MS→MO, SO→MS, MO→SS and SO→SS transition, resp. All kymographs are shown with a time resolution of 30 s. Time goes from left to right. The vertical scale bars represent 100 μm, the horizontal scale bars 60 min.

The oscillatory dynamics in SO and MO does not exhibit a regular period in many cases. This irregularity of repetitive protrusion events indicates rather a noisy excitable regime than regular oscillations in the strict sense of dynamical systems theory. We will see in section 2.2 that we find regular oscillations, noisy oscillations and a noisy excitable regime in our biophysical model.

Spread cells are symmetric and exhibit protrusions at both ends. Moving cells of course display protrusions at the front. However, we can identify additional back protrusions easily by the occurrence of negative back edge velocities in the oscillatory states as the cells in panels E and J of Fig. 1 exhibit. Hence, also moving oscillatory cells exhibit protrusions at the front and back. We cannot tell from panel F, whether a protrusion exists at the back of a cell in state MS while it moves steadily. If steady protrusions at the back exist, they are most likely shorter than front protrusions (see below). However, the emergence or extension of a back protrusion precedes the direction reversal in panel F by about 30 min. Hence, protrusions at front and back exist at the time of the transition which supports the idea of direction reversals being the result of the competition of front and back protrusions as we will see below.

### 2.2 Analysis of dynamic cell states

We analyse cell states on the basis of our biophysical model. Its components are explained in Fig. 2 and the equations are introduced in section 4.6 and the supplemental material. Eqs. 1-3 together describe the cell dynamics in terms of cell length and the retrograde flow friction coefficient *κ* determined by the number of bonds between the F-actin network and structures stationary in the lab frame of reference, i.e. the clutch state.

**Figure 2:**
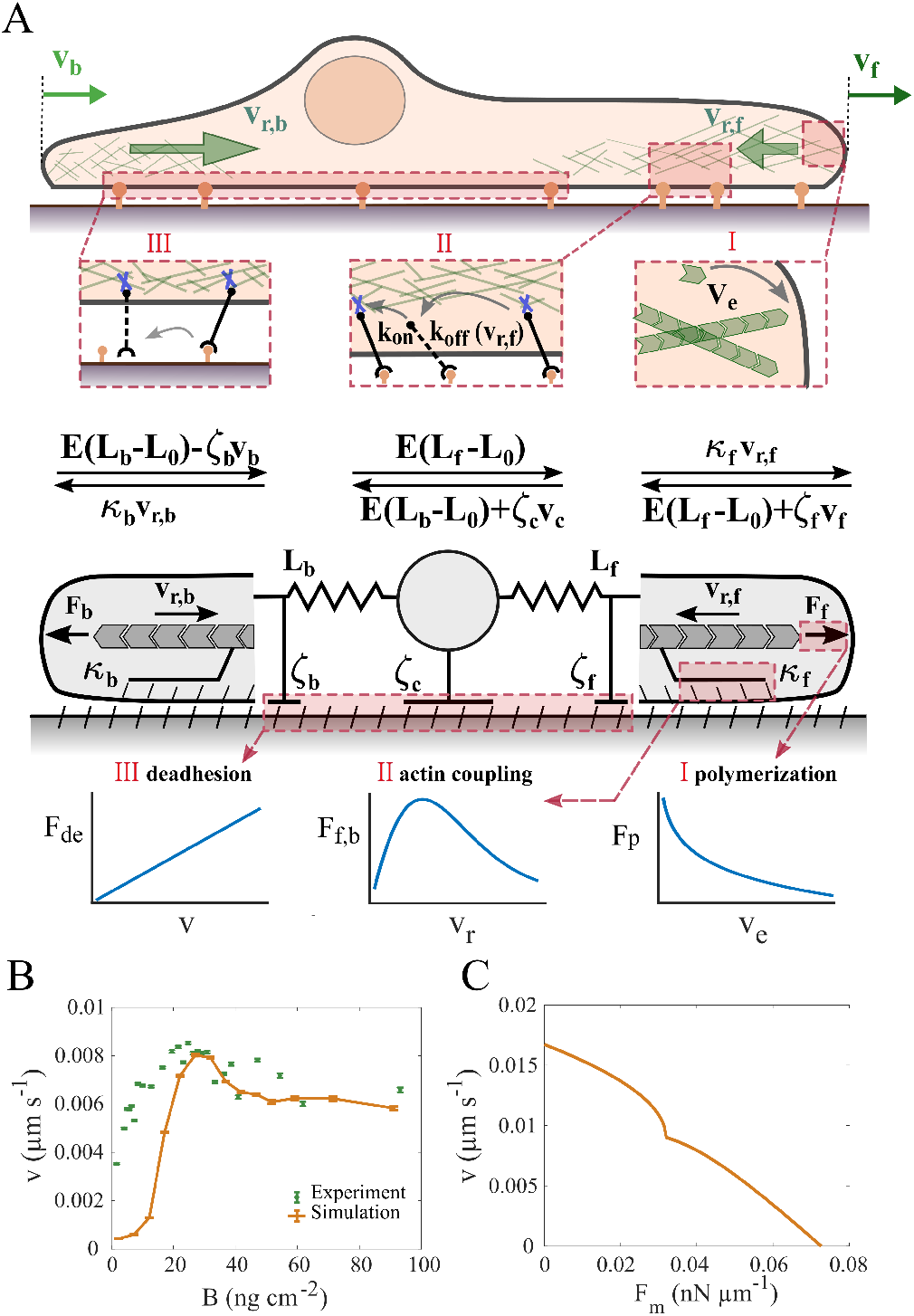
Model constituents. (A) Cartoon of a cell moving on a 1d Fibronectin lane (top), force balances (middle) and the mechanical components of the model (bottom). Front and back protrusion edges move with velocities *v_f_* and *v_b_*, resp. The F-actin networks flow with the retrograde flow rates *v_r,f_* and *v_r,b_*, resp. The forces *F_b_* = *κ_b_v_r,b_* and *F_f_* = *κ_f_ v_r,f_* arise from polymerisation of F-actin, act on the protrusion edge membrane and drive retrograde flow against the friction forces. The front and back edge membrane experience drag with the coefficients *ζ_f_* and *ζ_b_*, resp. Elastic forces *E*(*L_f_* – *L*_0_) and *E*(*L_b_* – *L*_0_) act between the cell body and the edges (equilibrium length *L*_0_). The balance of the elastic forces determines the motion of the cell body against the drag force *ζ_c_v_c_*. Bottom panels illustrate types of essential relations of the model. The de-attachment force of the back *F_de_* is linearly related to velocity. The friction force between F-actin network retrograde flow *v_r_* and stationary structures exhibits a maximum in its dependency on retrograde flow (clutch). The polymerisation force *F_p_* is logarithmically related to the network extension rate *v_e_* due to the force dependency of the polymerisation rate. **Two constitutive relations.** (B) The adhesion-velocity relation. Green dots represent experimental data, see table S1 - data set 7. Error bars represent the standard error of the mean. (C) The stationary force-velocity relation predicted by the model for *B*=45 ng cm^-2^ (see section S10 for details). It provides the cell velocity under constant application of the external force *F_m_* to the leading edge or cell body. Parameter set 1 from Table S2 for both panels.

Our model reproduces the adhesion velocity relation in agreement with experiments (Fig. 2B, section S5). This relation has been discussed in detail in Schreiber et al. [30] for cells with one protrusion in the direction of motion. The reproduction of this fundamental relation by the two-protrusion model supports our choice of modelling of the effect of Fibronectin signalling on friction and drag forces. The model predicts a stationary force-velocity relation of cell motion as shown in Fig. 2C. Our results are very similar to relations predicted earlier [30, 60]. All predictions agree in the point that this relation reflects the retrograde flow friction law [30, 60].

Analysis of cell states starts with appointing stretches of trajectories to one of the 4 dynamic states. The method of state classification is explained in section S2.1 and Fig. S1 in detail. It analyses cell behaviour described by kymographs. Deducing states from behaviour requires to define a minimal time of consistency. If the behaviour qualifies for this time as belonging to a specific state, we appoint this state to the cell. We have chosen 1h as this time (see section S2.1). We do not classify the state of simulated cells on the basis of the known dynamic regime of the model, but rather apply the same procedure to experimental and simulated data in order to maximise compatibility of outcome.

We draw our mechanistic conclusions on the basis of the agreement between model results and experiments. The model reproduces all 4 dynamic states of MDA-MB-231 cells (Fig. 3A). Both quantitative characteristics with regard to fraction of cells in the different states (Fig. 3B), oscillation period, oscillation amplitude and velocity, and qualitative ones like the back edge but not the front oscillating in the moving state are met by the model (Fig. 3A,E,F).

**Figure 3:**
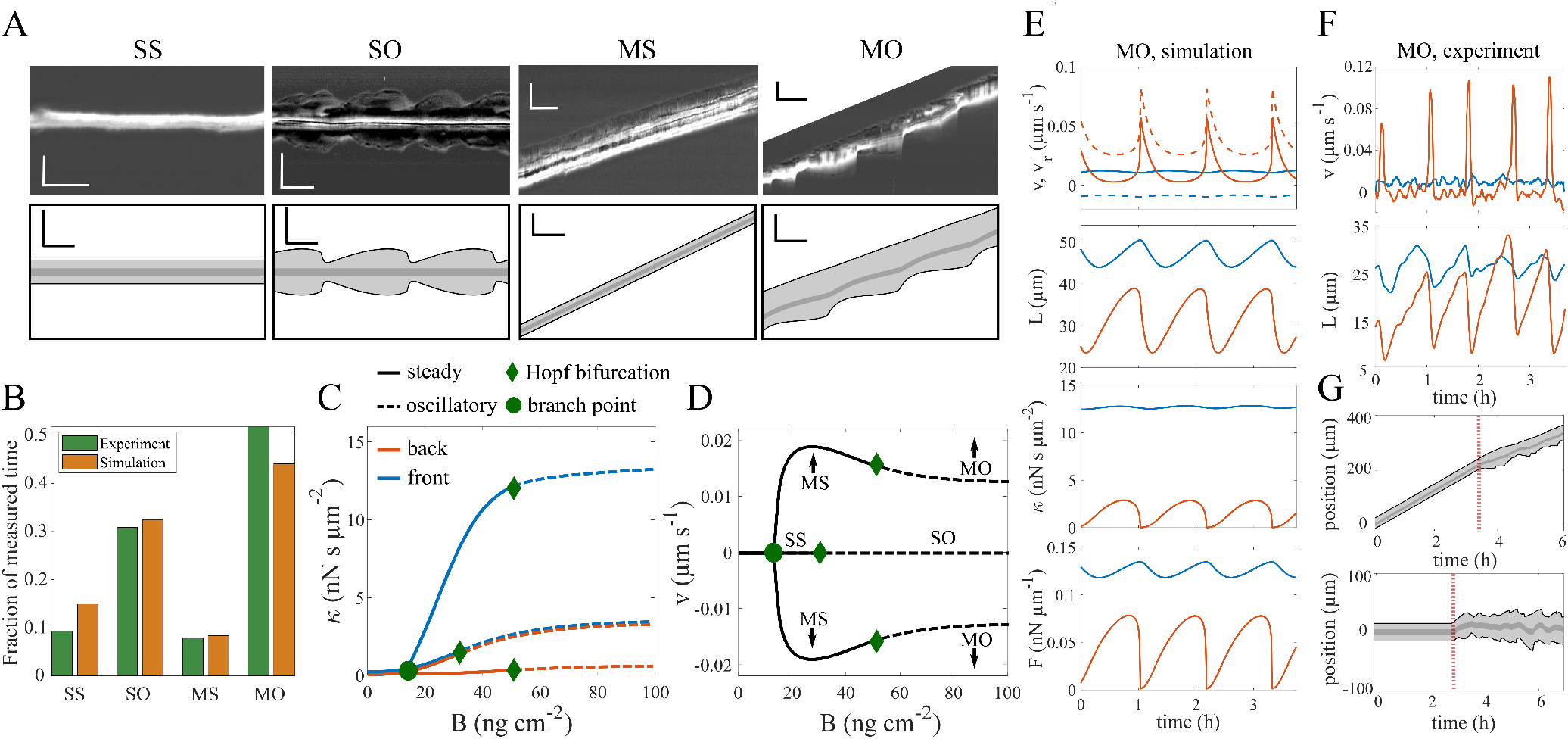
Analysis of dynamic cell states. They are either spread (SS, SO) or moving (MS, MO), and either steady (SS, MS) or oscillatory (SO, MO). (A) Upper panels - experiments, lower panels - simulations. Horizontal scale bars indicate 30 min, vertical scale bars 50 μm. The cut corner of the experimental MO state showed another cell. Simulations used *c*_3_=8.8 μms^-1^, SS, MS *B*=22 ng cm^-2^, SO *B*=36 ng cm^-2^, MO *B*=80 ng cm^-2^. (B) Fraction of cells in the cell states in experiment (2878 h of trajectories) and in simulations sampled from experiments on a range of Fibronectin concentrations (see section S1.2 for details). (C), (D) Cell states of the noise-free model illustrated by their value of the friction coefficient *κ* in the stationary state (C) and the cell velocity *v* (D) for Fibronectin concentration *B*. Only the state SS exists at low *B*. At the concentration denoted as branch point, moving states appear and coexist with the spread state. Oscillations start (dashed lines) at the Fibronectin concentrations marked as Hopf bifurcations. All moving states exist as up and down moving (D). (E) Simulated time course of the edge velocity v (full line), retrograde flow *v_r_* (dashed line), cell length *L,* friction coefficient *κ* and force on the edge membrane *F* in state MO (blue front, orange back). (F) Measured time course of edge velocity and cell length in state MO (blue front, orange back). (G) The steady states MS (upper panel) and SS (lower panel) are excitable. They behave steadily with noise free dynamics. Noise is switched on at the time marked by the red line. It causes behaviour very similar to noisy oscillations (Fig. S2). Parameters of all simulations are listed in set 1 in Table S2.

The force generation machinery at front and back protrusion work asymmetrically in the motile states MS and MO. This polarisation between protrusions does not require any signalling to be established. It is based completely on the mechanical properties of retrograde flow. Retrograde flow is always faster at the back than at the front, since it needs to keep up with cell motion (Figs. 2A, S7). Therefore, the biphasic friction force - retrograde flow velocity relation (Fig. 2, Eq. S19) entails that the strongest coupling between F-actin network and substrate via adhesion structures forms in the front protrusion. The value of *κ_f_* is always higher and forces are stronger in the front protrusion than in the back protrusion (Fig. 3E).

Driven stick-slip systems robustly generate oscillations [62, 63]. A stick-slip transition of the clutch is also here the core of the oscillation mechanism of the protrusions similar to earlier studies [42, 51, 52]. We describe it in detail in section S4. Whenever the forces driving retrograde flow drive it up to *v_cr_*, the flow slips causing the peaks in edge velocity and retrograde flow rate in Fig. 3E, F and a sudden drop of the friction coefficient and all forces. The recovery of *κ* afterwards is slow and takes the larger part of the period.

Retrograde flow velocity *v_r_* is equal to the network extension rate in the state SS. It is a tense state with two balanced opposing forces. This tense state is unstable against supercritical fluctuations in *κ*, which may arise from the concurrent snapping or formation of bonds between the F-actin network and stationary structures, and other perturbations as we will see below. The protrusion goes through one protrusion-retraction cycle upon a sufficiently strong perturbation. If these perturbations occur randomly, they are called noise. Chemical noise from random formation and breakage of bonds is omnipresent and represents relevant perturbations in systems as small as protrusions [64]. The relevance of noise for adhesion and retrograde flow dynamics has been demonstrated by a variety of studies [61, 65–68].

States in which small (but supercritical) perturbations may cause a large response are called excitable. Both the state SS and MS are excitable (Fig. 3G). The protrusion-retraction cycle after a perturbation in the excitable regime is very similar to the oscillation cycle of noisy oscillations (Fig. S2). Therefore, the permanent noise in the bonds between the F-actin network and adhesion structures may cause oscillation-like behaviour in the model cells even if they are in states SS and MS (Fig. 3G). Whether the noise is supercritical - causing oscillations - or not depends on the specific parameters of the cell and the noise amplitude. Thus, the states SS and MS with low noise amplitude coexist with oscillation-like states at higher noise amplitude.

The experimental and simulated oscillations shown in Fig. 3A are both rather smooth with subcritical noise, but noise may have a strong effect on state MO as we show in Fig. S2. It not only renders the time course irregular, but also substantially shortens the average period. Interestingly, we find examples for both smooth (Fig. 3A) and noisy oscillations (Fig. 1) in the experimental data indicating that noise amplitude is a cellular parameter and varies between individual cells. Accordingly, we have put the model noise in the clutch mechanism of retrograde flow, which is an intracellular process.

Figure 3C shows the existence of the dynamic states of the *noise-free* model cell in a systematic way for a range of Fibronectin coating densities. At coating densities below the value of *B* marked as branch point, only the spread state SS exists. Above it, spread and moving states coexist. Co-existence of several observable states for one set of parameters is multistability. At the Fibronectin density values marked as Hopf bifurcation points, oscillations start and we observe the states SO and MO. Between the branch point and the Hopf bifurcation of the spread branch, the spread state SS and moving state MS coexist. The Hopf bifurcation of the spread state occurs at smaller Fibronectin density (B) than the one of the moving state, and thus we find a range of coexistence of SO and MS. This order of bifurcation points is swapped at other parameter values and therefore SS can also coexist with MO (see Fig. S3, section S6). SO and MO coexist at large B-values.

An individual cell is described in the model by a set of parameter values. The population of cells in a given experiment represents many different parameter value sets due to cell-to-cell variability. Therefore, we may find all possible pairings of coexistence in a single experiment and both moving states can coexist with both spread states (Figs. 3C, S3).

In correspondence to its versatile experimental observation, multistability of the four morphodynamic states appears to be a very robust property of mathematical models of cells on 1d lanes including the clutch. Sens [52] and Ron et al. [51] also report very similar steady and oscillatory states (but not the excitable regime) and their coexistence in noise-free models. Hennig et al. report the states SS and MO and the irregularity of oscillations in a noisy stick-slip model [42].

### 2.3 Transitions between cell states

All possible transitions are illustrated in Fig. 4A. The fractions of transitions out of a given state are shown in Fig. 4B for the experimental data. The simulations in Fig. 4C show good agreement with the measurements. The same applies to the comparison of theory and experiment with Latrunculin A and Blebbistatin applied shown in Fig. S4.

**Figure 4:**
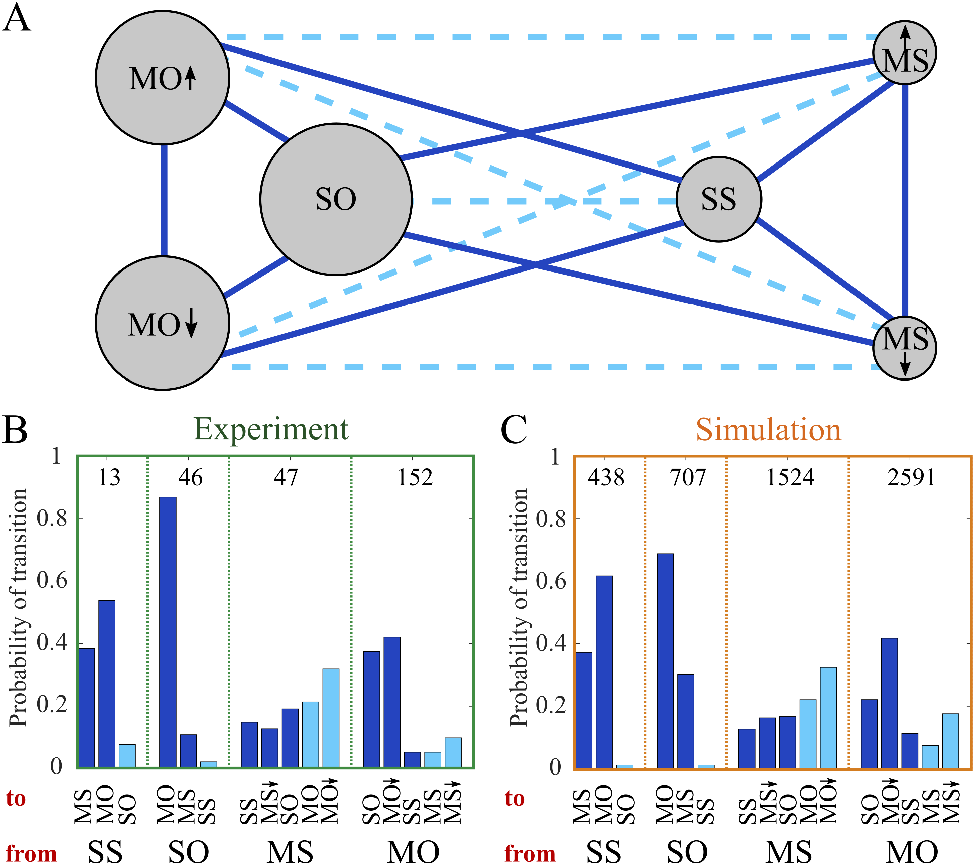
The transitions between the cell states. (A) The states MS moving steadily, MO moving oscillatory, SS spread steadily, SO spread oscillatory are introduced in Fig. 3 and the text. Arrows distinguish moving up and down. Solid lines mark transitions in the sense of dynamical systems theory of multistability, i.e., between states coexisting in the noise-free mathematical model. The state changes along dashed lines are explained in the text. (B), (C) Statistics of state transitions in experiments (B) and simulations (C). The bars belonging to a specific state listed at the bottom of the panel show the fraction of transitions out of this state to one of the other states. Transition types are colour-coded as in panel A. MO↓ and MS↓ indicate direction change in the transition. Analogous results with experimental data with Blebbistatin and Latrunculin A applied are provided in Fig. S4. Sample size is indicated by the numbers inside the chart. Parameters of all simulations are listed in set 1 Table S2.

Transitions between the dynamic states demonstrate that an individual cell can be in different states at given fixed conditions (or at the same parameter values in modelling terms). They are the experimental manifestation of multistability. The multistability in the biophysical model is shown in Figs. 3C, D, S3. Both spread states SS and SO can coexist with either MS or MO in the biophysical model. The upward-moving states coexist with their downward-moving analogues (Fig. 3C).

Transitions between these coexisting cell states are caused by noise in the adhesion variable in the biophysical model. For several reasons we assume that noise causes also the transitions in the MDA-MB-231 cells. Cells with their typical volume in the fl range are microscopic systems subject to thermal noise in many aspects of their behaviour [69–72]. Since adhesion sites are discrete spots, their length scales are even two orders of magnitude smaller than cell size rendering them even more susceptible to thermal noise. Our model results show that we can reach good agreement between theory and experiment by noise in the clutch mechanism of retrograde flow. In addition, transitions occur apparently spontaneously on homogeneous Fibronectin lanes without any obvious signalling event or stimulation.

We showed above that both motile states can coexist with both spread states in the model. However, SS does not coexist with SO, and neither MS coexists with MO in the bifurcation schemes in Figs. 3C, S3. How come we see transitions within these pairs of states in experimental and simulated data?

Oscillation-like protrusion events in the excitable regime occur randomly. Due to this randomness, cells may exhibit the characteristics of steady behaviour for a time sufficiently long to qualify as a state, and then switch to oscillation-like behaviour or vice versa. Thus, steady↔oscillatory-transitions come out of the state analysis of the data. However, they are not state transitions in the sense of dynamical systems theory for multistable systems but correspond to the onset or termination of random phases of consistent behaviour. Simulated and measured data behave very similar in regard to state classification and transitions - including steady↔oscillatory-transitions. Therefore, the mechanistic ideas formulated in the model reproduce the statistics of state transitions both in the sense of multistability and the statistics of phases of consistent behaviour of MDA-MB-231 cells.

The identification of steady↔oscillatory-transitions in simulated data implies that their occurrence in experimental data does not necessarily indicate parameter value changes. Changing parameter values might be an additional reason for these transitions in experiments, since cells constantly develop, and their cellular parameter values might change within the duration of an experiment of up to 48 h.

This section dealt with spontaneous state transitions. We present state transitions caused by Fibronectin steps in the supplementary material section S11. The biophysical model offers an explanation also for their characteristics.

### 2.4 Reversal of direction and the UCSP

Fig. 5A shows front and back edge velocity during a direction reversal averaged over many such events including both MS and MO reversals (see Fig. S7 for the retrograde flow, Fig. S2E for the network extension rate). The moment of reversal *t_rev_* is the time when the cell nucleus changes direction of motion. The back edge starts to slow down about 10 min prior to *t_rev_* and is already moving in the new direction (negative velocity) at the time of reversal. The front slows down after the back edge. It still moves in the old direction at *t_rev_* and reverses direction only a few minutes later. Finally, it collapses in an event appearing as a negative velocity peak in Fig. 5A. After the recovery of the protrusion, both edges move in the new direction with the same velocity. In agreement with this scenario, the likelihood of back protrusions increases before *t_rev_* and the frequency of occurrence of front collapses after *t_rev_*. Both experiments and simulations show the same scenario. A supercritical protrusion event at the back pulls sufficiently strong to collapse the front protrusion. The back protrusion event is caused by noise in the state MS. In the MO state, protrusions occur periodically but noise generates supercritical protrusion events. We discuss forces and retrograde flow during reversal events additionally in section S9 and Figs. S2, S7, S8.

**Figure 5:**
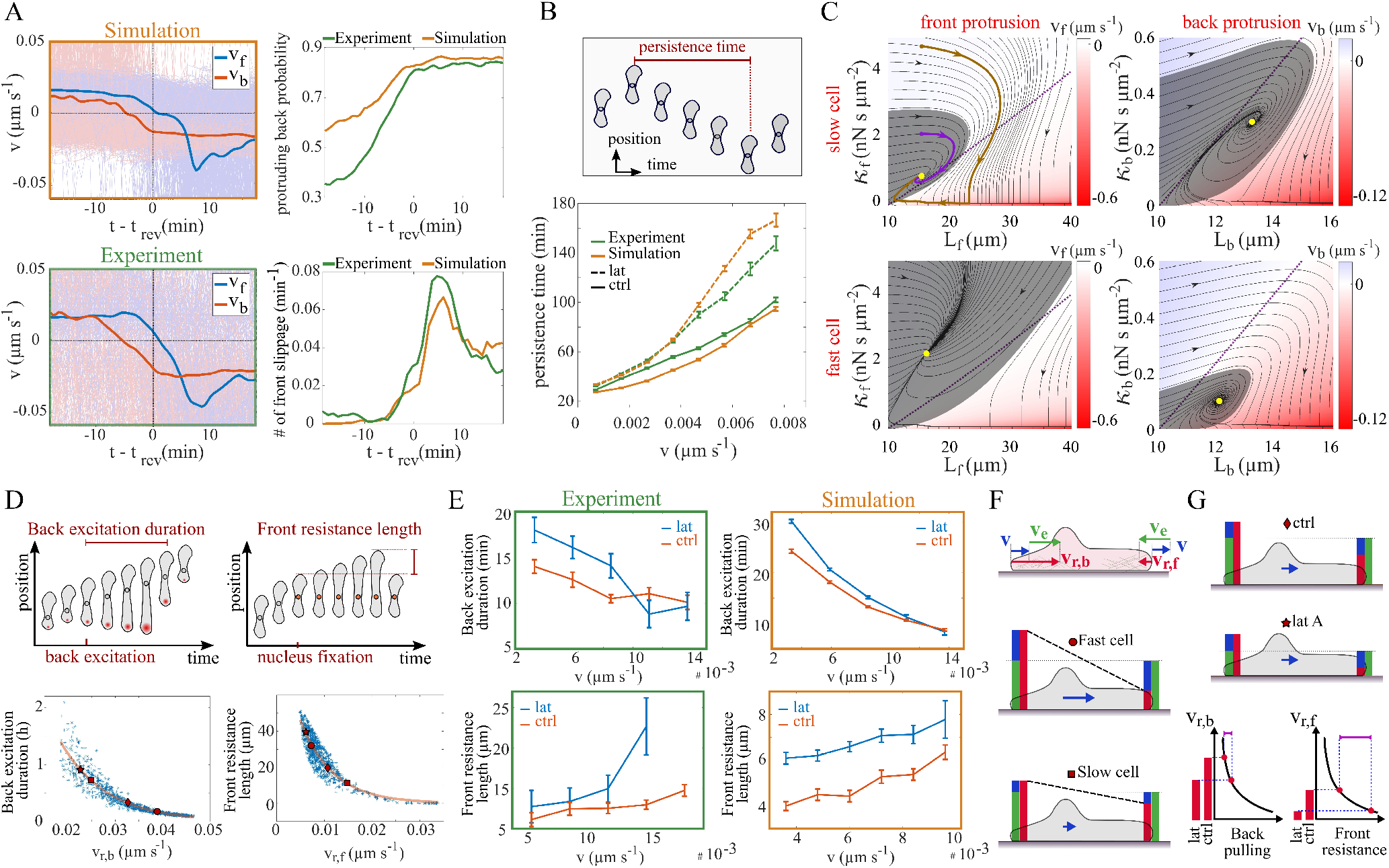
Direction reversal mechanism. (A) Front (*v_f_*) and back (*v_b_*) velocities during reversal transitions averaged over many tracks (thin lines, 2878 h experimental trajectories), including both MS and MO states. The cell nucleus changes direction at *t_rev_*. (B) The relation between cell velocity and persistence time in control cells (96577 h experimental trajectories) and with Latrunculin A applied (54809 h experimental trajectories). Latrunculin A application was modelled by decreasing the network extension rate 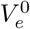 from 3.0 to 2.2 μms^-1^. (C) Basin of attraction of the steady protrusion state (grey area) and state trajectories (black lines) in *κ-L* plots. Trajectories outside the basin of attraction (brown) go to small values of the friction coefficient *κ* and then to small protrusion length *L* with fast retrograde flow, which is the collapse. Trajectories starting within the basin of attraction (purple) lead to the steady state (yellow dot) without collapse. B is 20 ng cm^-2^, velocities of fast and slow cells are 0.015 and 0.005 μms^-1^, respectively. (D) Cartoons illustrating definitions of duration of back excitation and front resistance length. Lower panels show simulations of these characteristics. Each dot marks the result of a simulation with parameter values randomly drawn from large parameter ranges (see section S1.2). (E) The relation between duration of back excitation and cell velocity, and front resistance length and cell velocity in experiments and simulations (2878 h control experimental trajectories, 2343 h Latrunculin A experiments). (F) Cartoon relations between velocities. Network extension rate (green) is the sum of cell velocity (blue) and retrograde flow velocity (red) in the front protrusion. Retrograde flow velocity is the sum of extension rate and cell velocity in the back protrusion, and therefore always faster than retrograde flow in the front protrusion. (G) Cartoon illustrating the effect of Latrunculin A. It reduces the network extension rate. Therefore, Latrunculin A treated cells have slower retrograde flow than control cells with the same cell velocity (colours as in F). This increases back pulling slightly and front resistance substantially (compare panel D) and thus renders the cells more persistent than control cells. Parameters of simulations in Table S2 (set 1 and 2 for control and Latrunculin conditions). Error bars represent the standard error of the mean.

Random direction reversal events are the reason why cells do not move permanently in their initial direction. Hence, the statistics of direction reversal events shapes the universal correlation between speed and persistence time (UCSP) which we show in Fig. 5B. We find an increase of persistence time with cell velocity (in agreement with earlier results [34]). Application of Latrunculin A increases persistence time. Latrunculin reduces the F-actin polymerisation rate and its application has been modelled accordingly by decreasing the force free polymerisation rate 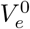 (set 2 in Table S2). We find good agreement between experiment and simulations both for control and Latrunculin A conditions, and also with Blebbistatin applied (see Fig. S5).

The velocity scenario in Fig. 5A, the observation that direction reversals happen only when protrusions at the front *and* back exist, and a strong robustness of the front protrusion against noise at the front all suggest direction reversal events to arise from competition between the front and back protrusion. As a first step in disentangling the competition mechanism we provide a picture of protrusion stability exemplified with the state MS. The values of the two dynamic variables friction coefficient *κ* and protrusion length L describe the state of the model protrusions. Upon a perturbation away from its steady state, a protrusion might just go back to the steady state or collapse (Figs. 5C, S6). The basin of attraction in a *κ-L* plot quantifies these two possibilities. If the protrusion is perturbed to a state within the basin of attraction, it relaxes back to the steady protrusion state. It collapses upon larger perturbations. Fig. 5C shows that the basin of attraction of the front protrusion of fast cells is larger than the one of slow cells and vice versa for the back protrusion. Hence, the front protrusion is more stable in fast cells than in slow ones, and the back protrusion is less stable in fast cells than in slow ones.

The pulling of the back protrusion on the nucleus and the front protrusion during the direction reversal scenario described above increases the elastic force acting on the front edge membrane. This force increase speeds retrograde flow up. If the back pulls strong enough, it drives retrograde flow velocity to the decreasing branch of the friction force - *v_r_* relation (see Fig. 2), and shifts the state of the front protrusion from the stationary state to a trajectory outside the basin of attraction, which entails the collapse. The collapse is a rapid decrease of *κ* due to breaking of bonds between the F-actin network and stationary structures followed by rapid shrinkage of protrusion length due to the elastic force as shown in Fig. 5C (see also description in section S8). Remarkably, the stochastic event inside the back protrusion, which in the end causes the front protrusion collapse and direction reversal, typically happens minutes before the moment of direction reversal.

The back pulls on the front by protruding (Fig. 5A), i.e., by going through an excitation in the state MS. The slower the retrograde flow in a back protrusion, the longer the excitation lasts (Fig. 5D). Whether it can collapse the front is determined by how long the front can resist the pulling. The slower the retrograde flow in the front protrusion, the longer it can resist. Symbols mark the retrograde flow values of typical fast and slow cells. Back excitations of fast cells are shorter and their fronts can resist longer than slow cells. These properties of front and back protrusion hold for a large parameter range, i.e. these relations between stability characteristics and retrograde flow are fundamental and robust features.

The front resistance length is the length the front edge moves after motion arrest of the cell body. Fig. 5E compares values for the duration of back excitation and front resistance length between our experimental data and simulations, and shows the qualitative agreement. Panel F summarizes our insights from the individual investigations. Fast cells have slow retrograde flow in the front protrusion and very fast retrograde flow in the back. This entails strong front protrusions and short excitations at the back and hence long persistence times. Slow cells have slower retrograde flow at the back and faster retrograde flow at the front, causing long back excitations and short front resistance length. They are therefore less persistent.

At a given velocity, Latrunculin A increases persistence (Fig. 5B). Application of Latrunculin reduces the network extension rate, and therefore reduces retrograde flow both at front and back compared to control. This increases the duration of back excitations but even more the resistance time and resistance length of the front and thus renders the cell more persistent (see Fig. 5B, E for experimental data and simulations). Fig. 5G summarises the Latrunculin action graphically. We conclude that reducing network extension rate increases persistence in agreement with our mechanistic ideas.

In summary, the protrusion competition mechanism based on elastic mechanical interaction between protrusions and cell body, non-linear friction of retrograde flow (clutch) and noise in the clutch mechanism offers an explanation for the UCSP. The faster cells move, the slower is retrograde flow in the front protrusion, the faster is it in the back protrusion and the more persistent the cell migrates.

## 3 Discussion

We analysed multistability of MDA-MB-231 cells on Fibronectin lanes, and found in our experiments coexistence of states with oscillatory or steady cell shape, spread or moving, moving up or down. We combined the experiments with quantitative theory, which suggests mechanistic ideas for several basic and general observations of mesenchymal motility comprising the biphasic adhesion-velocity relation, stationary force-velocity relation, UCSP, random migration and steady, oscillatory or excitable morphodynamics. Restricting cell motion to 1d made the relation between basic phenomena of cell motility very obvious. Random migration and the UCSP arise from random state transitions between the states ‘moving up’ and ‘moving down’, and the control of the parameters of the noisy clutch by integrin signalling generates the biphasic adhesion-velocity relation.

Our theory comprises three constituents, all of which are well established experimental observations. The first one is the force balance at the protrusion edge (Eqs. S2-S4). It establishes the link between polymerisation rate, cell velocity and retrograde flow velocity [19–21, 73–75]. The second constituent is the noisy non-linear friction between retrograde flow of the F-actin network and stationary structures (Eqs. S13, S14) known as clutch mechanism [61, 76–79]. It is crucial for oscillatory dynamics and multistability. Given a cell with protrusions at front and back and symmetry with regard to parameters of the protrusions, the clutch mechanism introduces the mechanical polarisation into a front protrusion with slow retrograde flow and a back protrusion with fast retrograde flow [52, 65]. We find that noise in the clutch mechanism due to random bonds between the F-actin network and stationary structures suffices to offer an explanation for state transitions and thus for the UCSP.

Multistability, oscillations and mechanical polarisation are all generated by the interaction of non-linear F-actin flow dynamics and membrane tension without any signalling processes. Signalling sets the parameters of this system and thus determines the dynamic regime (steady, excitable or oscillatory) and the cell velocity. A representation of the net effect of integrin signalling on drag and friction coefficients by Hill functions is the third constituent of the theory (Eqs. S17, S18). Integrin signalling together with friction and force balance determine the adhesion-velocity relation [30]. We know from Schreiber et al. [30], that a large part of cell type specific detail enters via this signalling constituent.

In order to support the idea that one mechanism explains the variety of observations, we followed the parameterization strategy to use one parameter set for all experiments with comparable conditions rather than fitting each experiment as good as possible. We were able to reach good quantitative agreement between simulations and experimental results with a single parameter set for control experiments and changes to parameter values describing drug applications corresponding to the known biochemical action of the drug. We could reach good agreement with regard to the types of states, the (temporal) characteristics of states, state fractions and transition probabilities, characteristics of reversal behaviour, the UCSP, protrusion stability, the adhesion-velocity relation and the behaviour on Fibronectin steps. The large variety of observations agreeing between theory and experiment is a major reason for our assumption that we assembled the model constituents dominating the observed behaviour, and that the mechanisms we suggest recapitulate the cellular processes. We review and discuss alternative mechanisms from literature in the supplement Section S13.

Our experimental data confirm the UCSP for MDA-MB-231 cells and thus add another cell type to the many listed in Maiuri et al. [34] obeying this relation. Fast cells are more persistent than slow ones. Maiuri et al. explain the UCSP by a mechanism centered around the network extension rate: the fast network extension in fast cells advects an F-actin-binding inhibitor of network growth away from the protrusion tip and thus renders random protrusion collapse unlikely. Maiuri et al. conclude that the faster the network extension rate the more persistent the cell moves [34]. However, the data in Maiuri et al. from mature bone marrow dendritic cells migrating in a confined environment exhibiting a positive correlation between network extension rate and persistence time do not obey the UCSP [34]. Furthermore, an endogenous F-actin binding inhibitor of network extension has not been identified. Contrary to these ideas, reduction of the network extension rate by Latrunculin entails an increase in persistence in our experiments and model. We found retrograde flow to be the most important indicator of stability (Fig. 5D). With our approach and our data, the noisy clutch is sufficient to explain the UCSP by spontaneous direction reversals based on protrusion competition.

The mechanism suggested by Maiuri et al. implies that increasing network extension rate necessarily accompanies increasing protrusion velocity, i.e. the authors require proportionality of the velocities *v_e_* = *avf* with *a* being constant across experimental conditions. Direct measurement of protrusion velocity and retrograde flow in keratocytes [33] and PtK1 cells [25], and the theoretical analysis of adhesion-velocity relations for keratocytes, PtK1 cells, CHO cells and MDA-MB-231 cells [30] come to the conclusion that this proportionality is violated. For example, increasing adhesion strength can reduce retrograde flow entailing increasing protrusion velocity with constant or even decreasing network extension rate. Data by Jurado et al. [76] and Vicente-Manzanares et al. [11] report the relation *v_e_* = *v_r,f_+v_f_* (our Eq. S5), which is different from proportionality since all three velocities depend on experimental conditions (see also Fig. S2F). Eq. S5 is obvious by geometrical reasons (Fig. 2) and also compatible with a variety of measured adhesion-velocity relations [30].

The stationary force-velocity relation of cell motion represents the cell response to external force and therefore a basic cell property. Due to technical problems of controlling either force or cell velocity to stay constant, it has not been measured yet, in difference to measurements and analyses of the dynamic relation, which allows both parameters to change during the experiment [19–21]. Our results are very similar to relations predicted earlier [30, 60]. All predictions agree in the point that the stationary relation reflects the retrograde flow friction law [30, 60].

We suggest as a main conclusion of our study that the basic phenomena multistability with its dynamic regimes, adhesion-velocity relation and UCSP can all be explained on the basis of the three model constituents force balance at the protrusion edges, noisy clutch and integrin signalling. They also entail a specific prediction for the stationary force-velocity relation of cell motion. All three model constituents are observations which have been established earlier and in other contexts. Our study connects them and thus reveals their explanatory power. The universality of the model constituents offers a simple explanation for the universality of the constitutive relations.

## 4 Methods

### 4.1 Cell culture

We cultured MDA-MB-231 breast cancer cells stably transduced with histone-2B mCherry (gift from Timo Betz, WWU Münster) in L15 medium with 2 mM Glutamax (Thermo Fisher Scientific, Waltham, MA USA) plus 10% fetal bovine serum (Thermo Fisher) at 37 °C. Cells were passaged every 2-3 days using Accutase (Thermo Fisher). For experiments about 5000 cells were seeded per dish. After 2-3 h cells adhered to the micropatterns and we exchanged the medium to L15 medium without phenol red. Then, we transferred the samples to the microscope and started measurements within 1-2 h. For inhibitor experiments 10 μM (+/-)-Blebbistatin (Cayman Chemical, Ann Arbor, MI USA), 100 nM Latrunculin A (Merck, Darmstadt, Germany), 0.25nM Calyculin A (Thermo Fisher) were added 2 h before the start of the experiment. As control we used DMSO (life technologies, Darmstadt, Germany) equal to the amount used for the dilution of the inhibitors.

### 4.2 Micropatterning

15 μm wide lanes which were homogeneously coated with Fibronectin (FN) were applied on an imaging dish with a polymer coverslip bottom (ibidi, Gräfelfing, Germany) using a micro-contact printing protocol. The production of the polydimethylsiloxane (PDMS) stamps and the subsequent printing has been described previously [80]. For all experiments a range of FN densities was covered.

### 4.3 Determination of Fibronectin densities

We determined the Fibronectin density via fluorescence intensity as described previously by us [30]. Lyophilised FN batches were resuspended and conjugated with Alexa Fluor 647 NHS ester (Thermo Fisher). We measured the concentration of FN in solution by optical absorption at 280 nm (Nanodrop, Thermo Fisher). The calibration factor that enables the conversion of fluorescence intensity to FN density was determined using microfluidic channel slides that were filled with a FN solution of known concentration.

### 4.4 Microscopy

We performed time-lapse imaging on an inverted fluorescence microscope (Nikon Eclipse Ti, Nikon, Tokyo, Japan) equipped with an XY-motorised stage and a heating chamber (Okolab, Pozzuoli, Italy) set to 37°C. Imaging was done using a 10x CFI Plan Fluor DL objective (Nikon), a CMOS camera (PCO edge 4.2, Excelitas PCO, Kelheim, Germany) and the acquisition software NIS Elements (Nikon). Before the start of the time-lapse measurement epifluorescence images of the FN patterns were taken. Then, phase contrast images and epi-fluorescence images of the cell nuclei were taken for 48 h at 10 min or 30 s intervals as indicated in the text.

### 4.5 Image analysis

Image analysis was performed using a combination of MATLAB R2020a (MathWorks, Natick, MA USA) scripts and FIJI (ImageJ) macros, based on previous work [30]. Fibronectin lanes are detected using a Hough transformation of the fluorescence signal of the labelled Fibronectin. The position of the nuclei is tracked by setting a threshold after applying a background correction and band-pass filter to the fluorescent images of the nuclei. The coordinates of the nuclei are converted such that the x-coordinates are parallel to the Fibronectin lanes. The position of the front and back of the cells is determined via kymographs that are created along the centre of the FN lane. Then the cell edge is manually segmented. Code used to generate results in the current study is available from the corresponding authors upon reasonable request.

### 4.6 Biophysical Model

We define the model constituents and their rate laws in this section. This will result in differential equations for the protrusion lengths *L_f,b_* and friction coefficients *κ_f,b_* for the front (f) and back (b) protrusion. Protrusion edge velocities *v_f,b_*, cell body velocity *v_c_*, retrograde flow velocities *v_r,f_, v_r,b_* and protrusion edge forces *F_f,b_* are determined by algebraic equations of *L_f,b_* and *κ_f,b_,* all listed in the supplement. Algebraic equations also relate the parameters of these dynamics to integrin signalling and the Fibronectin density.

Motivated by the observations presented in Fig. 1, we formulate a cell model with front and back protrusions moving on a 1d Fibronectin lane (Fig. 2A). The model is based in part on our previous work on steady cell motion in Schreiber et al. [30]. The main extensions of the model compared to Schreiber et al. [30] are the cell body, the back protrusion and the noisy clutch. The requirement for the noisy clutch from a modelling point of view follows from the dynamics we consider here, while we considered stationary properties in [30].

The force balances (see Eqs. S2-S4 and Fig. 2A) applying to the front and back edge and cell body have been established in a variety of studies [19–21, 60, 73–75, 81]. They comprise the drag forces resisting motion *ζ_f,b_v_f,b_* and *ζ_c_v_c_*, the retrograde flow friction force *κ_f,b_v_r,f,b_* and the elastic forces (Fig. 2A). The drag coefficients *ζ_f,b,c_* and the retrograde flow friction coefficients *κ_f,b_* are affected by adhesion and integrin signalling (Eqs. 2, 3, S17, S18). The cell body velocity *v_c_* is determined by the forces acting on it from front and back protrusions and the drag coefficient of the cell body *ζ_c_*.

We choose a linear dependency of the force between protrusion edges and cell body on the protrusion length (elastic force) based on the results in [30] (see Eqs. S2-S4) and [42]. We assume that the force is transmitted by membrane tension and consider as its most likely cause volume homeostasis in 1d [82].

The network extension rate *v_e_* is equal to the vectorial difference of the edge velocity and retrograde flow velocity. It is fixed by polymerisation, which is force dependent with the well known Arrhenius factor [73, 75, 83] (see Eqs. S5, S6).

The lengths of the protrusions *L_f,b_* are dynamic due to velocity differences between the edges and the cell body

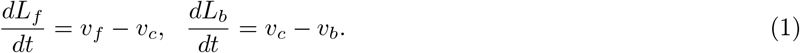

The noisy clutch has been reported in a variety of studies [61, 76–79, 84–86] and is due to the retrograde motion of the treadmilling F-actin network inside the protrusion [16, 87, 88]. This flow causes friction with all structures relative to which it moves, in particular also with stress fibres and the intracellular interface of adhesion sites [76, 89, 90]. The friction of the F-actin flow transmits the protrusion force to the substrate [21, 31, 76, 77, 89–92]. The value of the friction coefficient can be perceived as the state of the clutch, with large values corresponding to an engaged and low values to a disengaged state.

The relation between friction force and retrograde flow velocity exhibits increasing friction force at small velocities up to a critical velocity value *v_r,cr_* beyond which friction force decreases (Fig. 2A, Eq. S19). The force maximum entails stick-slip transitions of sudden acceleration at the critical velocity due to the decrease of the force resisting motion while the force driving motion is maintained. Stick-slip behaviour is a versatile phenomenon generating sound in bowed string instruments [62, 63], causing earthquakes [93] and wear in articular joints [94]. Recent theoretical studies suggested it to be relevant also for protrusion dynamics [42, 51, 52, 61, 65, 66] and polarisation [42, 52, 65].

The friction force is proportional to the number of transient bonds between the F-actin network and stationary structures in the protrusion. Its biphasic character is due to fast dissociation of these bonds at fast retrograde flow (the clutch disengages) [95]. They have to rebind to reach their equilibrium density after a high velocity phase. This motivates the *κ* dynamics adapted from [61]

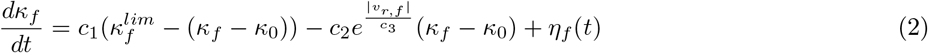

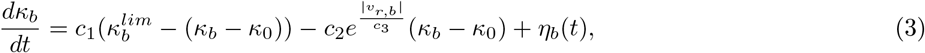

with an exponential acceleration of bond dissociation by retrograde flow velocity 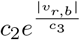 [95].

The maximum values 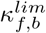 and *ζ_f,b_* exhibit a Hill-type relation to the Fibronectin substrate density *B_f,b,c_*, as specified by Eqs. S17, S18. This type of relation has been concluded from an earlier analysis of the adhesion-velocity relation [30]. As justified in detail in section 2.3, we assume that bond formation and breakage cause some noise on top of the deterministic dynamics, and added the noise terms *η_f_*(*t*) and *η_b_*(*t*) (see also Eq. S16). The stationary states of Eqs. 2, 3 represent the biphasic friction force - retrograde flow velocity relation Eq. S19. The detailed equations are provided in the supplementary material. The parameter values of the model are listed in Table S2.

## Supporting information

data_set_1_ctrl_30s

data_set_2_lat_30s

data_set_3_blebb_30s

data_set_4_ctrl_10min

data_set_5_lat_10min

data_set_6_blebb_10min

data_set_7_untreated_10min

## 5 Data Availability

Experimental data supporting the findings of this manuscript are available from the corresponding authors upon reasonable request. Source data of cell trajectories is provided as CSV files in the SI. A description of the data sets is available at SI Appendix, section S1.

## 6 Acknowledgments

We thank K. Rottner and T. Stradal, HZI Braunschweig, for stimulating discussions. This work was supported by the Deutsche Forschungsgemeinschaft (DFG201269156, SFB1032, J.C.J.H., C.S. and J.O.R.).

## 7 Author contributions

B.A. executed the theory and analysed data, J.C.J.H., C.S. executed experiments and analysed data, M.F. designed and supervised theoretical research, J.O.R. designed and supervised experimental research, J.C.J.H., B.A., J.O.R., M.F. wrote the paper. All authors have read and approved the published manuscript.

## 9 Additional information

### 9.1 Supplementary information

The online version contains supplementary material.

