## Supplementary material for "On multistability and constitutive relations of motion of MDA-MB-231 cells on Fibronectin lanes": SI Appendix

#### S1 Data sets and simulations

##### S1.1 Data sets

Experimental data was recorded in two different temporal resolutions, 30 s and 10 min, respectively. Data with high temporal resolution allows the study of the dynamics of the cell edges. But the number of cell tracks per experiment is limited by the fact that only a limited number of field of views can be covered during a time step of 30 s. The lower temporal resolution of 10 min enabled the acquisition of much bigger data sets which in turn facilitate the study of characteristics such as persistence time as defined below. In the data sets with 10 min temporal resolution, about 22900 single-cell tracks that are moving on homogeneous Fibronectin lanes with densities in the range of 0-120 ngcm$^{-2}$ were analysed. The data sets with 30 s temporal resolution comprise of about 400 single-cell tracks (about 6400 h) on homogeneous Fibronectin lanes with densities in the range of 0-40 ngcm$^{-2}$. Data sets 1-3 consist of data from 4 independent experiments acquired with 30 s temporal resolution and data sets 4-7 of data from 6 independent experiments with a temporal resolution of 10 min.

For high temporal resolution data sets 1-3 the first column contains the cell track ID, the second the time in the lab frame (min), the third the position of the centre of nucleus along the FN lane (μm), the fourth the median FN density along the FN lane of the cell (ng cm$^{-2}$), the fifth the position of the cell's upper edge along the lane (μm), the sixth the position of the cell's lower edge (μm), the seventh the position of the nucleus' upper edge, the eighth the position of the nucleus' lower edge, and the ninth the state change points. For the simulation data sets the tenth and eleventh columns contain the retrograde flow velocity at the upper and lower protrusions respectively, and the twelfth column contains the critical retrograde velocity of each cell. For low temporal resolution data sets 4-7 the first column contains the cell track ID, the second the time in the lab frame (min), the third the position of the nucleus along the FN lane (μm) and the fourth the median FN density along the FN lane of the cell (ng cm$^{-2}$).

| Name of data set | Temporal resolution | Treatment | Number of cell tracks | Total trajectory time | Figures |
|---|---|---|---|---|---|
| 1_ctrl_30s | 30 s | control | 221 | 2878 h | 1,3-5 |
| 2_lat_30s | 30 s | Latrunculin A | 127 | 2343 h | 5, S4 |
| 3_blebb_30s | 30 s | Blebbistatin | 65 | 1165 h | S4 |
| 4_ctrl_10min | 10 min | control | 9497 | 96577 h | 5 |
| 5_lat_10min | 10 min | Latrunculin A | 3368 | 54809 h | 5 |
| 6_blebb_10min | 10 min | Blebbistatin | 3728 | 47638 h | S5 |
| 7_untreated_10min | 10 min | untreated | 6261 | 65378 h | 2 |
| stepped lanes | 10 min | untreated | 6158 | 76759 h | S11 |

Table S1: List of data sets containing experimental cell trajectories.

Data for cell trajectories on stepped FN lanes is taken from [1]. This data set comprises of about 6200 single-cell tracks. For a list of all experimental data sets, see Table S1.

#### S1.2 Simulations

All simulations were performed in MATLAB (Mathworks). We have 2 experimental control data sets (set 1 and the step experiments set 4 in Table S2) with the experiments done in different months. Set 1 comprises experiments on homogeneous lanes and has been used for the data in Figs. 3, 4 and 5. Set 4 are the experiments on stepped Fibronectin lanes in Fig. S11. Parameter values obtained from the fits vary slightly between the two sets most likely due to day-to-day variability of cell behaviour (Table S2).

From a modelling point of view, drug treatment appears in the model as parameter change resulting from fits to experimental data with the corresponding drug applied. Latrunculin reduces the actin polymerisation rate [2]. Very much in agreement with this known action of Latrunculin, a reduction of the polymerisation rate $V_e^0$ from 0.03 $\mu$ms$^{-1}$ in set 1 control simulations to 0.022 $\mu$ms$^{-1}$ in set 2 Latrunculin simulations was sufficient to fit the Latrunculin data in Fig. 5.

The myosin inhibition upon Blebbistatin application affects the integrin signaling pathway, which is manifested by the variation of $\kappa$ and $\zeta$. In an earlier study on the adhesion-velocity relation fitting data of 4 different cell types with very good agreement to a model with one protrusion only, we also allowed for contractile action of myosin [1]. It turned out to be negligible in all 4 cell types including MDA-MB-231 cells. Also now, changes in the values of the parameters $\kappa^{max}$ and $\zeta^{max}$ were sufficient to simulate the effect of Blebbistatin treatment (set 3 in Table S2). Our finding here, that the action of Blebbistatin can be fit by modifying the representation of the integrin signalling pathway only, very much confirms the earlier results, and are in agreement with results reported by Hennig et al. [3] for RPE1 cells and NIH-3T3 fibroblasts. Hence, we find the effect of myosin as part of the integrin signalling pathway to be more relevant than as contractile driver of retrograde flow in MDA-MB-231 cells.

We simulated a total number of approximately 6100 cell tracks with a length of 15 hours each. That includes 2500 cell tracks for the control condition, 2800 cell tracks for the Latrunculin condition, and 800 cell tracks for the Blebbistatin condition. Simulations start with a randomly chosen initial state of the cell. The Fibronectin densities on lanes are homogeneous in simulation sets 1, 2, and 3. Cell tracks in each experimental data set comprise a range of Fibronectin densities. We used the same values of Fibronectin densities in the simulations. All analyses in Figs. 3-5 are carried out by averaging over the Fibronectin density values with an ensemble of simulations with the same weight of the Fibronectin densities as in the experiments.

In case of the simulations for the experiments on stepped lanes, we reproduced the exact densities on the individual steps of the individual lanes and averaged within the ensemble of simulations with the same rules as on homogeneous lanes. For the stepped lanes we simulated a total number of approximately 1200 cell tracks.

Cell to cell variability is an important factor in the experiments affecting the population's behaviour. Therefore, we included the cell variability in the simulations by parameter variability. The model parameters in each simulation set are chosen according to Table S2, with an allowed variability of $\pm 5\%$ to mimic cell variability in experiments.

The simulations in Fig. 5D were performed with an allowed variability of $\pm 50\%$. For simplicity of the analysis, we only consider the cell tracks that are in the MS state when the noise is switched off. In the back excitation simulations (left panels in Fig. 5D), cells move (MS state) for 3 h without noise in the system. Then we apply noise to the back protrusion (Eq. S14). A back excitation is defined as a peak of cell length with at least 3 µmamplitude. The back excitation duration for each cell track is determined by averaging the duration of all back excitation events. The back retrograde flow for each cell track is calculated from the noise-free steady state.

In the front resistance simulations (right panels in Fig. 5D), noise is switched off throughout the simulations. We arrest the cell body motion instantaneously after 3 h of steady state motion. The front edge continues to move, which stretches the front protrusion. The front resistance length is defined as the difference between the maximum cell length before collapsing and the noise-free steady state cell length. The front retrograde flow for each cell track is calculated from the noise-free steady state before the nucleus fixing.

### S2 Data analysis

We used the kymographs of single-cell tracks to analyse the cell states. To exclude the interactions of cells, we terminated a single-cell kymograph when it got too close to a neighbouring cell. Then we found the position of the cell edges by segmenting manually the kymographs in FIJI (ImageJ) using a self-made macro. The same manual segmentation process was used for the nucleus, resulting in the data of the position of the nucleus edges (Fig. S1A).

#### S2.1 Classification of cell states

Our classification method described below analyses cell behaviour obtained from kymographs. We used stretches of trajectories excluding interactions between cells which entails stretches shorter than the duration of the experiment. Deducing states from behaviour requires the definition of a minimal time of consistency $t_c$. If the behaviour during this time qualifies as belonging to a specific state, we appoint this state to the cell. We chose 1h as this time, which is

| Parameter | Set 1: control | Set 2: Latrunculin | Set 3: Blebbistatin | Set 4: stepped lanes | Units |
|---|---|---|---|---|---|
| Figures | Figs. 2, 3, 4, 5, S2, S3, S6, S7, S8, S9 | Figs. 5, S4 | Figs. S4, S5 | Figs. S10, S11 | |
| $E$ | 3e−3 | 3e−3 | 3e−3 | 3e−3 | nNμm$^{-1}$ |
| $L_0$ | 10 | 10 | 10 | 10 | μm |
| $V_e^0$ | 3e−2 | 2.2e−2 | 3e−2 | 3e−2 | μms$^{-1}$ |
| $k^-$ | 5e−3 | 5e−3 | 5e−3 | 3e−3 | μms$^{-1}$ |
| $c_1$ | 1.5e−4 | 1.5e−4 | 1.5e−4 | 2e−4 | s$^{-1}$ |
| $c_2$ | 7.5e−5 | 7.5e−5 | 7.5e−5 | 1e−4 | s$^{-1}$ |
| $c_3$ | 7.8e−3 | 7.8e−3 | 7.8e−3 | 8e−3 | μms$^{-1}$ |
| $\kappa^{max}$ | 35 | 35 | 20 | 30 | nNsμm$^{-2}$ |
| $K_\kappa$ | 35 | 35 | 35 | 20 | ngcm$^{-2}$ |
| $n_\kappa$ | 3 | 3 | 3 | 3 | |
| $\kappa_0$ | 1e−2 | 1e−2 | 1e−2 | 1e−1 | nNsμm$^{-2}$ |
| $\zeta^{max}$ | 1.4 | 1.4 | 1.2 | 2.4 | nNsμm$^{-2}$ |
| $K_\zeta$ | 50 | 50 | 50 | 50 | ngcm$^{-2}$ |
| $n_\zeta$ | 4 | 4 | 4 | 4 | |
| $b$ | 3 | 3 | 3 | 2 | |
| $\zeta_0$ | 1e−1 | 1e−1 | 1e−1 | 2e−1 | nNsμm$^{-2}$ |
| $\alpha$ | 4e−2 | 4e−2 | 4e−2 | 4e−2 | nN$^{-2}$s$^{-2}$μm$^4$ |

Table S2: Parameters used for simulations. Parameter set 1 is used for simulations in control condition, set 2 for Latrunculin, and set 3 for Blebbistatin conditions. Simulations with sets 1, 2, and 3 are all carried out on homogeneous Fibronectin lanes. Parameter set 4 is used in the simulations on stepped Fibronectin lanes.

sufficiently larger than typical oscillation periods of about 15 min. and allowed to distinguish steady and oscillatory states. Detection of transitions requires stretches of tracks of at least $2t_c$ in order to identify the state before the transition and the state after it as different. Since transitions happen rarely just in the middle of a trajectory, stretches of several times $2t_c$ are required for state transition analysis. That requirement limits the length of $t_c$ from above.

###### S2.1.1 State change points

We split the single-cell tracks into different states. For this reason, a method is employed which uses an iterative change-point analysis based on cumulative sum (CUSUM) statistics similar to [4] to find the transition points between the two states of a cell. This algorithm is able to detect the time points with a fundamental change of motility trend. The period between two subsequent change points is defined as an episode of the cell with a specific state. Any episode lasting shorter than 1 h is disregarded and appended to the preceding episode. The detected state change points are shown in Fig. S1B for a cell track.

###### S2.1.2 State identification

We employed two criteria to determine the closest match to the states SS, SO, MS, and MO during an episode of the cell. First, to assess whether the cell is moving or spread, we compare the average velocity of the cell during

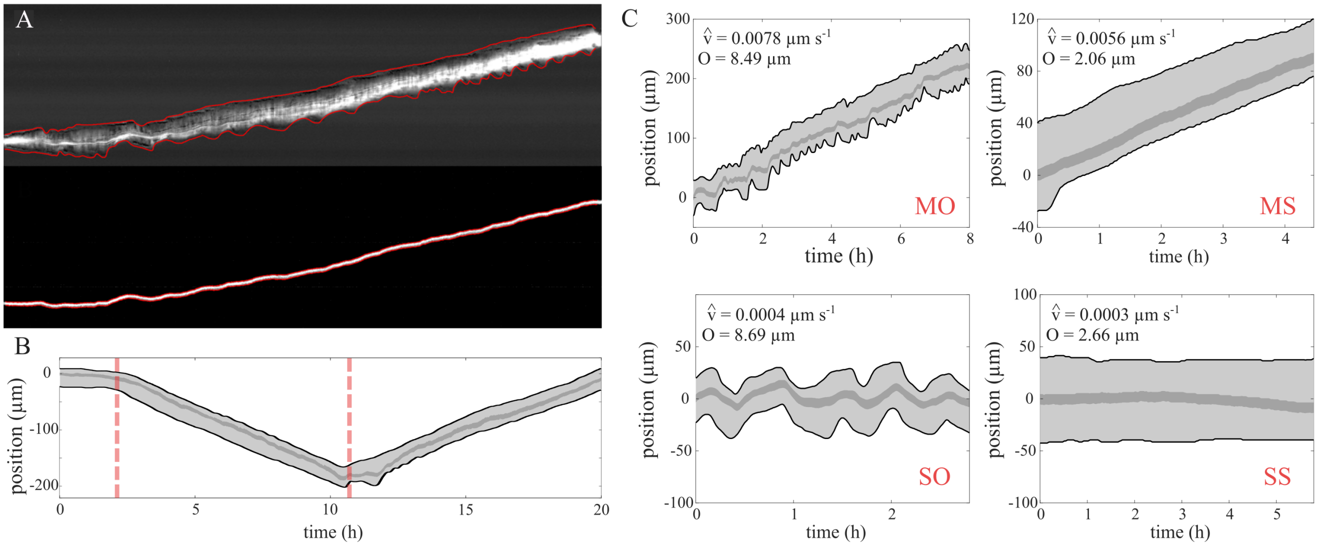


Figure S1: Classification of cell states. (A) Manual segmentation of the cell (upper panel) and nucleus kymographs (lower panel). The red lines show the segmented outlines of the kymographs. (B) State change points of a cell track. The dashed red lines indicate the time frames of a state change of the cell. (C) Examples of identified states. The values of the average velocity $\hat{v}$ and oscillation metric $O$ shown in each panel determine the state of the cell.

the episode with a critical speed (0.002 $\mu\mathrm{ms}^{-1}$ ). Cells with an average velocity smaller than this critical speed are considered spread.

The many cases of out-of-phase relations of the observed oscillations entail not only length dynamics but also motion of the nucleus. It turned out, that we get the most reliable classification when including both the length dynamics and the nuclear position dynamics. In the first step, we remove the small fluctuations of the length and cell body position. These fluctuations do not represent the oscillations caused by the mechanisms discussed in this study. Similarly, prolonged slow length changes are also not caused by the oscillation. They can be attributed to very slow cell processes that change cell properties like membrane tension and elasticity on long time scales. Long-term trends of cell body position are also related to the general motion of the cell rather than nucleus movements due to the cell oscillations. So, we apply a band-pass filter to the length and cell body position data to remove the tiny fluctuations and long trends. We chose the cutoff frequencies of 1 and 6 $\mathrm{h}^{-1}$ to remove the variations with time scales shorter than 10 min and longer than 1 h.

This filtered data contains the variations of length and cell body position in the time scales relevant to the oscillations caused by the competition of protrusions. For an oscillatory state, filtered length and/or cell body position vary a lot, while they stay almost constant in time for a steady state. Thus, the summation of the average absolute deviation of the filtered length and the average absolute deviation of the filtered cell body position is an indicator of the degree of oscillations.

$$O = \frac{1}{n}\sum_{i=1}^{n}(|L_{f,i}| + |X_{f,i}|)] \qquad \text{(S1)}$$

$O$ is the oscillation metric, $n$ is the number of time frames of the state, $L_{f,i}$ and $X_{f,i}$ are the band-passed filtered length and cell body position of the cell at time frame $i$, respectively. We then compare this oscillation metric at each cell episode with a critical value (5 µm). We find this critical value to classify cell states optimally by manual inspection of the cell episodes and their respective values of the oscillation metric. Fig. S1C shows examples of 4 cell episodes with their respective average velocity and oscillation metric values and thus their identified state.

##### S2.2 Motion metrics

Persistence time is defined as the average time during which a cell body maintains its moving direction on the 1D Fibronectin lanes. We calculate persistence using the low temporal resolution data with time frame intervals of 10 min. We only consider the moving episodes lasting at least two time frames (20 min). With this cutoff, small fluctuations do not affect the persistence calculation.

To measure the duration of back excitations (Fig. 5E), we only look into the moving states of the cells. Front and back are then defined according to the direction of movement. We then find the episode of back protruding. The duration of these episodes is averaged over a moving state of a cell. The average velocity of cell body during the moving state is then used to plot Fig. 5E.

We identify collapse events of the front by negative front velocity and a minimal retraction length of 2 µm. The length that the front protrusion shrinks during the complete retraction episode is interpreted as the resistance length in the analysis of the experimental data. We then find the average velocity of the front edge during the protruding episode right before the collapse. This velocity and the shrinkage length during retractions are used to find the experimental relation between front resistance length and velocity (Fig. 5E).

#### S3 Model

Our model describes a cell with front and back protrusion moving on a 1d cell track. Eqs. S2-S4 are the force balance of the mechanical system shown in Fig. 2 for the front(S2), cell body (S3) and back (S4).

$$F_f - E(L_f - L_0) - \zeta_f v_f = 0 \tag{S2}$$

$$E(L_f - L_0) - E(L_b - L_0) - \zeta_c v_c = 0 \tag{S3}$$

$$-F_b + E(L_b - L_0) - \zeta_b v_b = 0 \tag{S4}$$

The force-length relation of the coupling between front and back is well described by an elastic spring [1]. This coupling force can be membrane tension or any length-dependent cytoskeletal force. The front and back edge motion experience a drag with the coefficients $\zeta_f$ and $\zeta_b$, resp. The cell body velocity is determined by the forces acting on it from front and back protrusion and the drag coefficient of the cell body $\zeta_c$. $F_f$ and $F_b$ in the above equations are defined such that positive sign indicates pushing the membrane out.

The vectorial difference of the edge velocity and retrograde flow velocity $v_r$ is the network extension rate [5]. It is fixed by polymerisation, which is force dependent. A few publications motivated us to include this force dependency of polymerisation into our model. Renkawitz et al. [6], Fig. 2A of Maiuri et al. [7] (see also our Fig. S2), and McGrath et al. [8] showed that this dependency is possibly relevant. Bieling et al. measured an exponential dependency like an Arrhenius factor for the single filament polymerisation rate at small forces [9]. Network effects like changes in filament density due to branching modify the exponential single-filament dependency at about one quarter of the force-free network extension rate towards less steep decrease [9]. Bieling et al. used a system without retrograde flow. Systems with retrograde flow exhibit much weaker and/or modified density changes, since increasing force also speeds up retrograde flow [10, 6, 11]. Another network effect is in some cases binding of filaments to the obstacle surface enabling them to exert pulling forces [12, 13, 14, 15, 6]. The force dependence has been worked out for a variety of filament-obstacle interaction potentials by Motahari and Carlsson [16]. It leads to the well known Arrhenius factor for constant pushing forces [17, 13], which is also a good approximation with a variety of interaction potentials [16]. On that basis and on the basis of the low force data in ref. [9] we choose:

$$v_{r,f} + v_f = V_e^0 exp(\frac{-aF_f}{N}) - k^- \tag{S5}$$

$$v_{r,b} - v_b = V_e^0 exp(\frac{-aF_b}{N}) - k^-. \tag{S6}$$

Here, the factor $a = gd/k_BT$ subsumes a factor arising from the average over filament orientation in the network, the length $d$=2.7 nm added by an actin monomer to the filament and the thermal energy $k_BT$. $N$ is the number of filaments per edge contour length, and $k^-$ is the depolymerisation rate. We have chosen the value N=248 µm$^{-1}$ in all simulations similar to Schreiber et al. [1]. That value entails $a/N$=1 µm nN$^{-1}$. An evaluation in retrospect on the basis of our results presented in Fig. S2 showed that the force feedback to network extension rate is not essential in our mechanisms .

The force balancing the drag forces at the protrusion edge is the force $\kappa_f v_{r,f}$ driving retrograde flow [18]. With Eqs. S2, S5 and $F_f = \kappa_f v_{r,f}$ we find

$$v_f = \frac{N_f}{a\zeta_f} W_0\left(\frac{V_e^0 a\kappa_f\zeta_f}{N_f(\zeta_f+\kappa_f)} exp\left(\frac{a\kappa_f(k^-\zeta_f - E(L_f-L_0))}{N_f(\zeta_f+\kappa_f)}\right)\right) - \frac{E(L_f-L_0)+k^-\kappa_f}{\zeta_f+\kappa_f}, \tag{S7}$$

and analogously for the back velocity

$$v_b = -\frac{N_b}{a\zeta_b} W_0\left(\frac{V_e^0 a\kappa_b\zeta_b}{N_b(\zeta_b+\kappa_b)} exp\left(\frac{a\kappa_b(k^-\zeta_b - E(L_b-L_0))}{N_b(\zeta_b+\kappa_b)}\right)\right) + \frac{E(L_b-L_0)+k^-\kappa_b}{\zeta_b+\kappa_b} \tag{S8}$$

The retrograde flow velocities $v_{r,f}$ and $v_{r,b}$ can also be written as a function of $L$ and $\kappa$

$$v_{r,f}(\kappa_f, L_f) = \frac{1}{\kappa_f} W_0\left(\frac{V_e^0\kappa_f\zeta_f}{(\zeta_f+\kappa_f)} exp\left(\frac{\kappa_f(k^-\zeta_f - E(L_f-L_0))}{(\zeta_f+\kappa_f)}\right)\right) - \frac{-E(L_f-L_0)+k^-\zeta_f}{\zeta_f+\kappa_f} \tag{S9}$$

$$v_{r,b}(\kappa_b, L_b) = \frac{1}{\kappa_b} W_0\left(\frac{V_e^0\kappa_b\zeta_b}{(\zeta_b+\kappa_b)} exp\left(\frac{\kappa_b(k^-\zeta_b - E(L_b-L_0))}{(\zeta_b+\kappa_b)}\right)\right) - \frac{-E(L_b-L_0)+k^-\zeta_b}{\zeta_b+\kappa_b} \tag{S10}$$

For simplicity we have chosen the reasonable value $N_f = N_b = 248\ \mu\text{m}^{-1}$ entailing $\frac{a}{N_f} = \frac{a}{N_b} = 1\ \mu\text{m/nN}$ (see also [1]). The velocity of the cell body obeys Eq. S3 as, $v_c = \frac{E(L_f-L_b)}{\zeta_c}$. The length of the protrusions $L_{f,b}$ changes according to the velocity difference between edge and cell body

$$\dot{L_f} = v_f - v_c \tag{S11}$$

$$\dot{L_b} = v_c - v_b \tag{S12}$$

The friction resisting retrograde flow is proportional to the number of transient bonds between the F-actin network and stationary structures in the protrusion. The biphasic relation between retrograde flow velocity and friction forces [5, 19, 20, 21, 22] is caused by dissociation of these bonds at high values of the velocity [23]. They have to reform to reach their equilibrium density after a high velocity phase. This motivates the bond dynamics [24]

$$\frac{d\kappa_f}{dt} = c_1(\kappa_f^{lim} - (\kappa_f - \kappa_0)) - c_2 e^{\frac{|v_{r,f}|}{c_3}}(\kappa_f - \kappa_0) + \eta_f(t), \tag{S13}$$

$$\frac{d\kappa_b}{dt} = c_1(\kappa_b^{lim} - (\kappa_b - \kappa_0)) - c_2 e^{\frac{|v_{r,b}|}{c_3}}(\kappa_b - \kappa_0) + \eta_b(t), \tag{S14}$$

143

$$\langle \eta_{b,f}(t) \rangle = 0 \tag{S15}$$

$$\langle \eta_{b,f}(t)\eta_{b,f}(t') \rangle = \frac{c_1(\kappa_{b,f}^{lim} - \kappa_{b,f}) + c_2 e^{\frac{|v_{r,b,f}|}{c_3}}\kappa_{b,f}}{\alpha\kappa_{b,f}^{lim}}\delta(t-t') \tag{S16}$$

with an exponential acceleration of bond dissociation by retrograde flow velocity [23]. $\kappa_{f,b}^{lim}$ is the upper limit of $\kappa$ which is determined by the substrate adhesion strength. The terms $\eta_{b,f}$ add Gaussian white noise, arising from the random formation and breaking of bonds of the F-actin network with stationary structures. Our recent study showed that the description of the equilibrium values of $\kappa_{f,b}$ and $\zeta_{f,b}$ by a Hill-type relation with the Fibronectin substrate coating density $B_{f,b,c}$ provided a quantitative description of the adhesion-velocity relation in a single-protrusion model [1]. These findings enter the $\kappa_{f,b}$-dynamics by

$$\kappa_f^{lim} = \kappa_0 + \frac{\kappa^{max} B_f^{n_\kappa}}{K_\kappa^{n_\kappa} + B_f^{n_\kappa}}, \qquad \kappa_b^{lim} = \kappa_0 + \frac{\kappa^{max} B_b^{n_\kappa}}{K_\kappa^{n_\kappa} + B_b^{n_\kappa}}. \tag{S17}$$

The drag coefficients $\zeta$ at different regions of the cell are:

$$\zeta_f = \zeta_0 + \frac{\zeta^{max} B_f^{n_\zeta}}{K_\zeta^{n_\zeta} + B_f^{n_\zeta}}, \qquad \zeta_b = \zeta_0 + \frac{\zeta^{max} B_b^{n_\zeta}}{K_\zeta^{n_\zeta} + B_b^{n_\zeta}}, \qquad \zeta_c = b\left(\zeta_0 + \frac{\zeta^{max} B_c^{n_\zeta}}{K_\zeta^{n_\zeta} + B_c^{n_\zeta}}\right) \tag{S18}$$

$\zeta_0$ and $\kappa_0$ are the base level of the drag coefficient and friction coefficient. The factor $b$ describes the contribution of the cell body to the cell drag compared to the protrusion. Note that relations for the number of bonds and drag at front and back are symmetrical. Eqs. S11-S12, S13, and S14 form a 4th order dynamical system, which determines the cell motility behaviour. We find that our model can reproduce 4 different states of a cell.

Finally, the stationary states of Eqs. S13-S14 exhibit the biphasic friction force - retrograde flow velocity relation discussed in the introduction,

$$F_{f,b} = v_{r,f,b}\left(\kappa_0 + \frac{\kappa^{lim}}{1 + \frac{c_2}{c_1} e^{\frac{|v_{r,f,b}|}{c_3}}}\right), \tag{S19}$$

with the maximum force at $v_{r,cr}$ (see Fig. S7).

### S4 The oscillation mechanism

We describe here the oscillations of the state SO. The mechanism in the state MO is essentially the same. The spread states are symmetric and it is therefore sufficient to consider one protrusion (without variable indices $f, b$) as we do in Fig. S2. We start the description of the oscillation cycle in the phase when $\kappa$ starts to decrease. Retrograde flow $v_r$ increases driven by the force $F_f = E(L - L_0) + \zeta_f v_f$ in this phase (Fig. S2). The increase of $v_r$ causes further decrease of $\kappa$ due to bond rupture and it continues to fall. Since retrograde flow becomes faster, leading edge velocity $v$ slows down - the growing elastic force due to growing length $L$ brakes additionally. The drag force $\zeta v$ vanishes with decreasing edge velocity $v$. At some point, $\kappa$ is so small that the friction force cannot balance the elastic force anymore. The F-actin network slips with a large peak of $v_r$. That causes a further and faster decrease of $\kappa$. The leading edge follows the network with a negative velocity peak, which rapidly decreases $L$ and the elastic force and $F$ collapse. Retrograde flow has lost its driving force at this moment and $v_r$ and $v$ slow down immediately. Since the elastic force is very small now, the drive for retrograde flow is small and $v_r$ drops very low. The drop of $v_r$ has $\kappa$ start to rise again, making $v_r$ slow down further. The approximate conservation of the network extension rate has the protrusion velocity go up as $v_r$ decreases (Eq. S5), till the elastic force limits it. That is the time of minimal $v_r$ and maximal $v$. From that moment on, since $v>0$ still holds, $L$ and the elastic force continue to grow and speed up retrograde flow. The share of retrograde flow in the network extension rate increases again. The growth of $\kappa$ stops

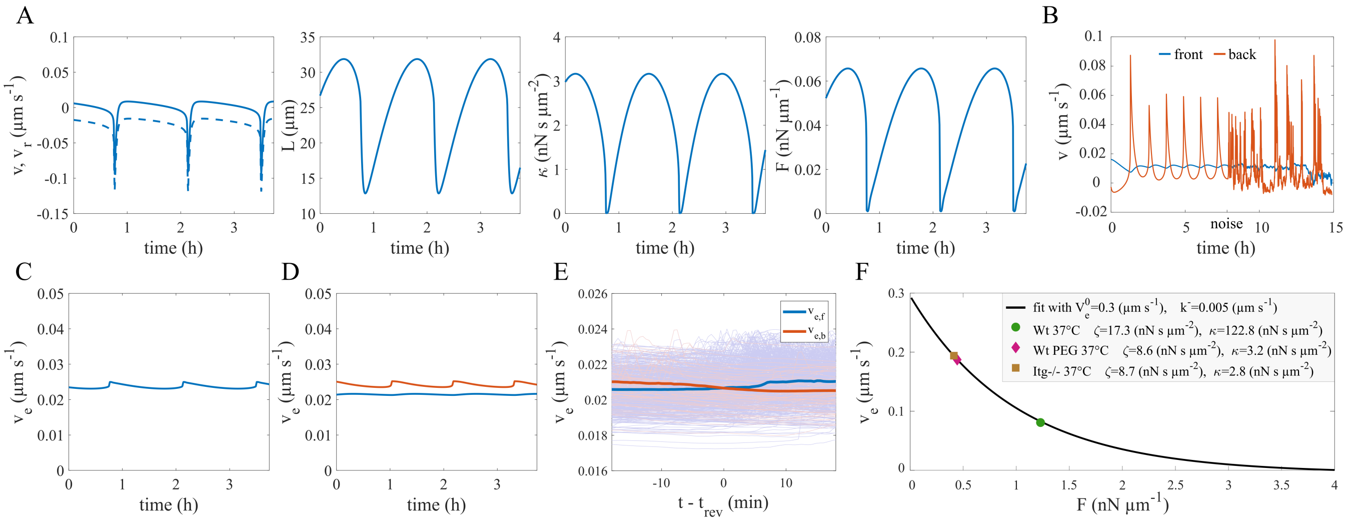


Figure S2: (A) Simulated time course of the edge velocity $v$ (full line), retrograde flow $v_r$ (dashed line), cell length $L$, friction coefficient $\kappa$ and force on the edge membrane $F$ in state SO. Front and back protrusion behave identical in deterministic simulations of this state. (B) The simulation in the state MO starts deterministic and then switches on the noise (blue front, orange back). Remarkably, the average frequency of the noisy oscillations is higher than the deterministic one. The deterministic oscillations are most similar to the smooth experimental example in Fig. 3A, the noisy oscillations are more similar to the noisy oscillations in Fig. 1. (C-E) The time course of the network extension rate in state SO (C) and MO (D) and as average over many simulations during a direction reversal event (E). Its changes are very small. Consequently, the force dependency of polymerisation is not an essential feedback in our oscillation and direction reversal mechanisms. The results in A-C and D are obtained from simulations of SO and MO states in Fig. 3, resp. Simulations in A-D use parameter values of set 1 listed in Table S2 with $c_3 = 8.8$ μms$^{-1}$. The Fibronectin density B in panels A-C and D are 36 ng cm$^{-2}$ and 80 ng cm$^{-2}$, resp. The results in E are obtained from simulations similar to Fig. 5A with the parameter values of set 1 listed in Table S2. (F) Maiuri et al. [7] present in their Fig. 2 data not obeying the UCSP. We consider here the data from this figure measured at 37°C. The green dot shows control data ($v = 0.071$ μms$^{-1}$, $v_r = 0.01$ μms$^{-1}$). The other two dots show conditions with substantially increased retrograde flow velocity due to reduced interaction between retrograde flow, actin cortex and substrate by either lack of Fibronectin ligand (PEG, purple, $v = 0.051$ μms$^{-1}$, $v_r = 0.138$ μms$^{-1}$) or integrin knock out (Itg-/-, brown, $v = 0.047$ μms$^{-1}$, $v_r = 0.146$ μms$^{-1}$). We have fit these 3 data points to Eq. 4 of Schreiber et al. [1] providing the relation between friction coefficient $\kappa$, cell drag coefficient $\zeta$ and cell velocity, and to the Eq. S5 for the dependency of the network extension rate on force (black line). The force-free network extension rate $V_e^0$, $\kappa$ and $\zeta$ are the fit parameters. In agreement with the experimental conditions, both $\kappa$ and $\zeta$ are substantially reduced with PEG and Itg-/-. The force-feedback to network extension rate is relevant here, since the network extension rate changes by about a factor 3 compared to control while the value of $V_e^0$ is the same for all three conditions. The relevance of this feedback has also been suggested by Renkawitz et al. [6].

and turns again into a decline since $v_r$ increases. This closes the loop.

### S5 Biphasic adhesion-velocity relation

The adhesion velocity relation has been measured in untreated cells (data set 7_untreated_10min). We used the control parameter value set 1 (Table S2) for these simulations. We averaged the velocity over the whole population in the experiments at a given Fibronectin density. We find a biphasic relation with maximal cell velocity at intermediate Fibronectin densities, as in Schreiber et al. [1]. We find that our simulations for control condition (parameter value set 1 in Table S2) can reproduce the biphasic adhesion-velocity-relation observed in the experiments (Fig. 2B). Similar to [1], we find saturation of the velocity at large Fibronectin densities in simulations and experiments. This shows that our model with two protrusions also reproduces the biphasic adhesion-velocity relation which is a general experimental observation with many cell types.

### S6 Remarks on other possibilities for the coexistence of states

In Fig. 3C, we showed the coexistence of SO and MS states in a range of Fibronectin densities (see also Fig. 4A). This is caused by the fact that the Hopf bifurcation of the moving state occurs at higher Fibronectin density (B) than the Hopf bifurcation on the spread branch. The order of Hopf bifurcation points on the moving and spread branches of Fig. 3C may be different for another cell described by a slightly different set of parameter values. Fig. S3A shows a case in which the Hopf bifurcation of the spread state occurs at higher Fibronectin density, and thus SS and MO states coexist in a range of B. We used the parameter values of set 1 in Table S2 and only increased the parameter $c_3$ to 0.009 $\mu\text{ms}^{-1}$ for this case. We also find that SS and MO or SO and MS states can coexist in broader ranges of B. Fig. S3 B and C show two cases with wide ranges of SS-MO and SO-MS coexistence, respectively. In these cases, the Hopf bifurcation point on either the spread state branch (panel B) or the moving state branch (panel C) does not appear in the range of B between 0 and 100 ng $\text{cm}^{-2}$. Possible transitions between the states are illustrated in the lower panels of Fig. S3 for each case. SO and MO states are not present in the noise-free model in panels B and C, respectively. However, the steady states can still be excitable in the noisy model and exhibit oscillation-like excitation events. Fig. S3 shows that steady states (SS and MS) can be found also at very high Fibronectin densities with some cells described by the corresponding parameter values.

In summary, due to cell-to-cell variability within a given experiment, we find a larger variety of coexistence options than indicated by a single bifurcation scheme.

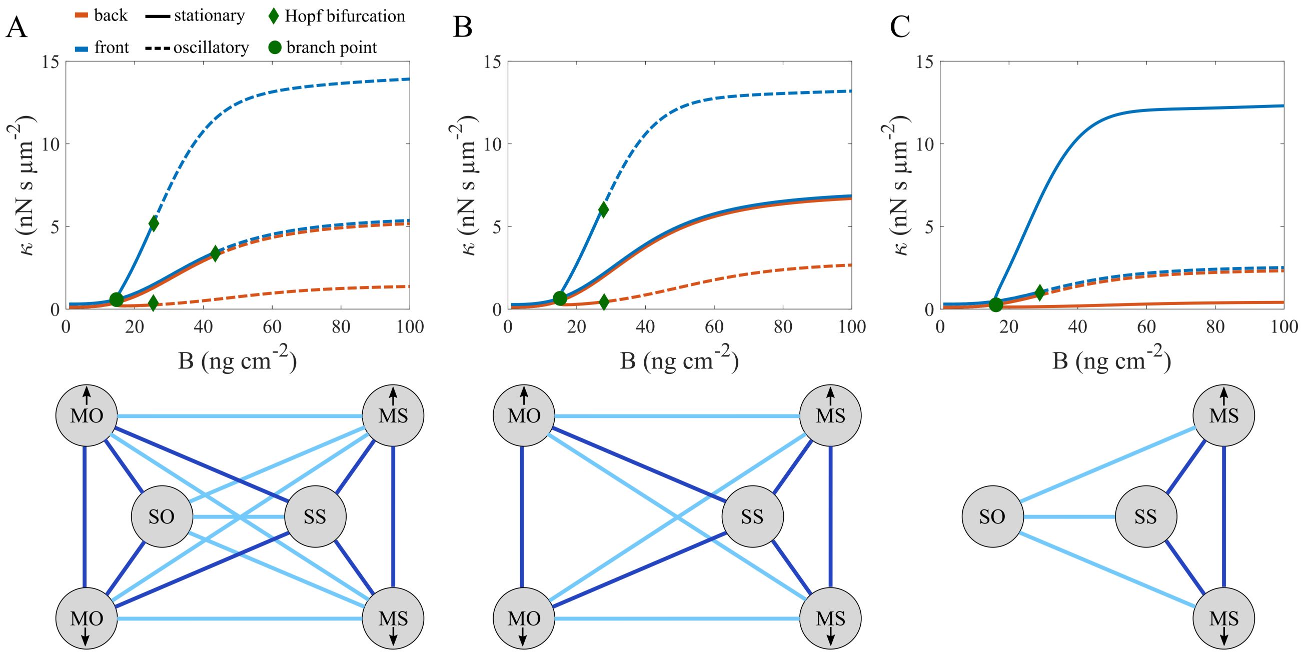


Figure S3: The dynamic cell states and transitions between them with 3 different parameter value sets. The upper row shows the cell states of the model illustrated by their value of the friction coefficient $\kappa$ for a range of Fibronectin density B. Dashed lines show the unstable stationary state in the oscillatory regime. The lower row shows the possible transitions between cell states. Dark blue lines show the transitions between states coexisting in the noise-free mathematical model. The parameter values are taken from set 1, Table S2, with (A) $c_3 = 0.009$ μms$^{-1}$, (B) $c_3 = 0.01$ μms$^{-1}$, and (C) $c_3 = 0.007$ μms$^{-1}$. The SO and MO states do not appear in (B) and (C), respectively, within the studied range of Fibronectin density B.

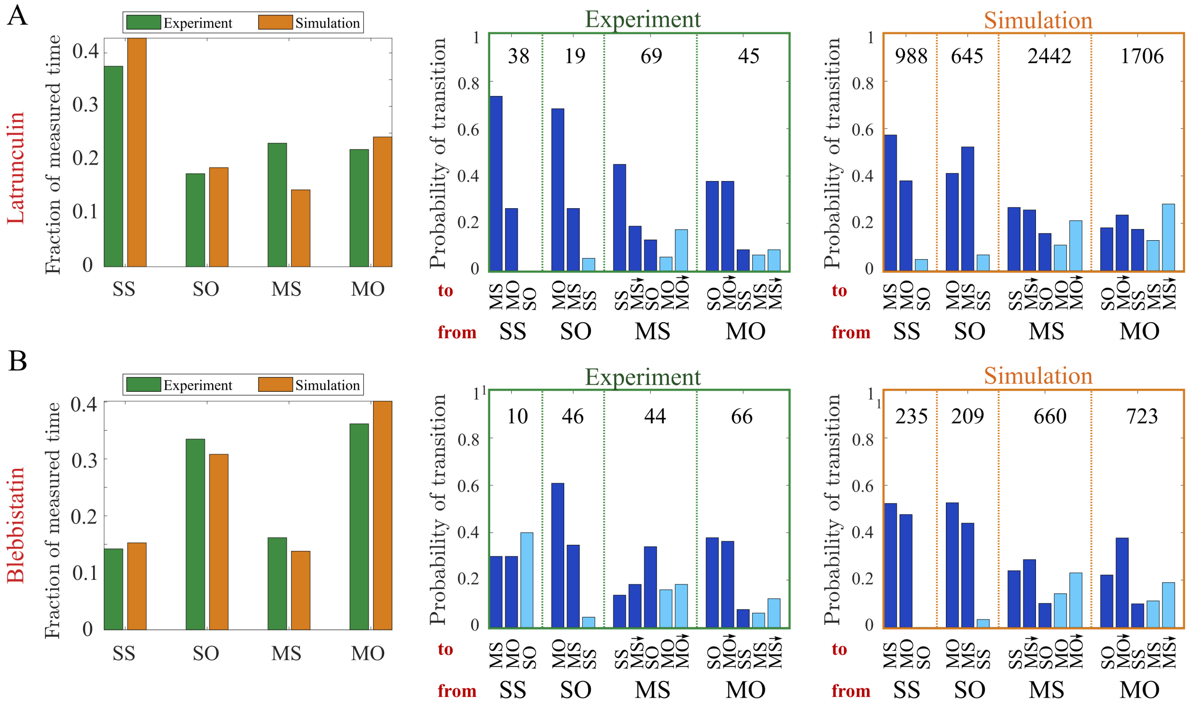


Figure S4: Analysis of dynamic cell states and transitions in Latrunculin A (A) and Blebbistatin (B) treated cells. Left column: Fraction of cells in the dynamic cell states in experiments and simulations in a range of Fibronectin densities. Middle column: Statistics of state transitions in experiments. Right column: Statistics of state transitions in simulations. The parameter values of data sets 2 and 3 (Table S2) are used for Latrunculin A and Blebbistatin simulations, respectively. Sample size is indicated by the numbers inside the chart.

### S7 Remarks on the effect of the drug treatments on states and transitions and persistence

Fig. S4A and B show the probability of states and transitions for Latrunculin and Blebbistatin-treated cells. We find that the application of Latrunculin increases the fraction of cells in the spread states compared to control condition (Fig. 3B). The fraction of steady states relative to the oscillatory states is also increased. The simulations in Fig. S4A show good agreement with the measurements. Only the network extension rate $V_e^0$ is decreased relative to control parameter values in Latrunculin simulations (Table. S2) which corresponds to the effect of the drug on the system. Thus, lower probability of oscillatory and moving states is a result of lower network extension rate. Also the probabilities of state transitions in the simulations show good agreement with the measurements in the Latrunculin-treated cells. Fig. S4B shows that the fraction of states in Blebbistatin-treated cells is almost similar to the control condition.

Blebbistatin application does not change the UCSP qualitatively. We find that persistence time increases with cell velocity also in the Blebbistatin-treated cells (Fig. S5). Application of Blebbistatin increases the persistence time compared to control, but not as much as the Latrunculin application. The simulations of the Blebbistatin condition show good agreement with the measurements (Fig. S5).

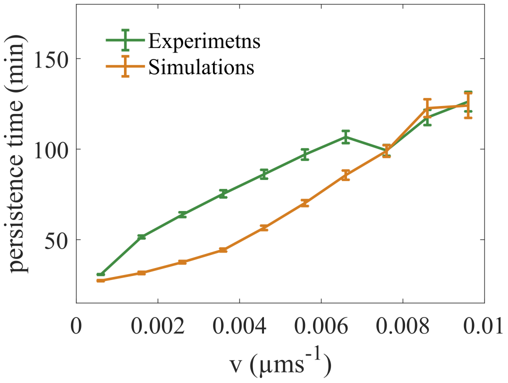


Figure S5: The relation between cell velocity and persistence time with Blebbistatin applied (modeled by parameter values of set 3 in Table S2).

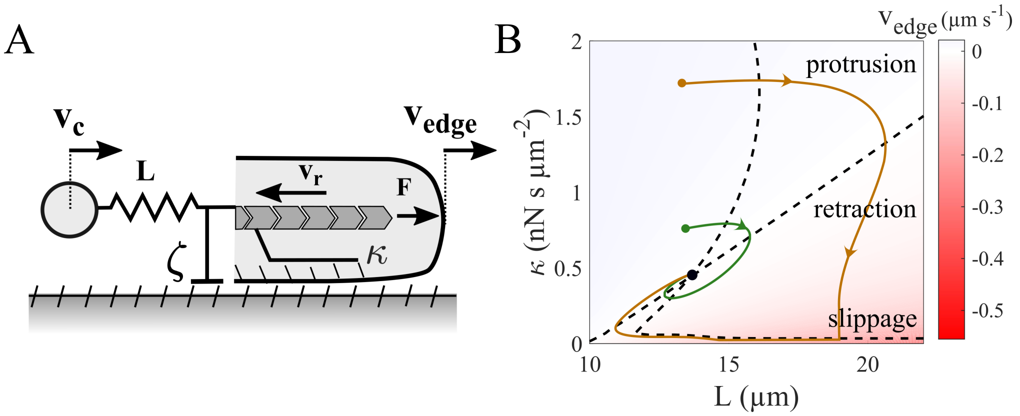


Figure S6: (A) Schematic illustration of the model for one protrusion. Here ($v_c$) is a model parameter which comprises the effects from the other cell parts on the protrusion. (B) The phase plane of the spread state case ($v_c = 0$). Two black dashed lines show the nullclines of the system. Green and orange lines are two example trajectories in the phase plane; one goes smoothly to the stable state (green), and one goes to the stable state through a slippage retraction (orange). The background colour indicates the velocity of the cell edge. Above and below the diagonal line the edge is in protrusion and retraction, respectively. The bottom right corner of the plane is associated with slippage phenotype. The parameter values of set 1 (Table S2) are used with B = 20 ng cm$^{-2}$ for the phase plane in this figure.

### S8 Excitability and stability of a protrusion in the cellular states SS and MS as seen by dynamic systems theory

Dynamic systems theory uses the terms 'state' and 'phase space' to characterise system behaviour rather than plots of dynamic variables in dependence on time. A state of a system is completely described by the value of its dynamic variables[1]. A dynamic variable obeys a differential equation fixing its dynamics. A dynamical system has as many dynamic variables as ordinary differential equations describing it. Each dynamic variable defines one dimension (or coordinate axis) of the phase space. A point in the phase space corresponds to a state of the system. We have 4

[1]Our use of the term 'cell state' for SS, MS, SO, MO deviates from this definition. The four cell states correspond to dynamic regimes in terms of dynamical systems theory.

dynamic variables: $\kappa_f$, $\kappa_b$, $L_f$ and $L_b$.

We consider steady protrusions in the cellular state SS and MS in this section. If the protrusions are very weakly coupled, it suffices to consider a single protrusion, which has the 2 dynamic variables $\kappa$ and $L$. With this assumption, the effect of the other cell parts (the cell body and the protrusion at the other end of the cell) on the protrusion is included in the parameter $v_c$ (Fig. S6A). Specifying the model to this situation leads to

$$\frac{d\kappa}{dt} = c_1(\kappa^{lim} - (\kappa - \kappa_0)) - c_2 e^{\frac{|v_r|}{c_3}}(\kappa - \kappa_0) \tag{S20}$$

$$\frac{dL}{dt} = v - v_c = \frac{1}{\zeta} W_0\left(\frac{V_e^0 \kappa\zeta}{(\zeta+\kappa)} e^{\frac{\kappa(k^- \zeta - E(L-L_0))}{(\zeta+\kappa)}}\right) - \frac{E(L-L_0) + k^-\kappa}{\zeta+\kappa} - v_c \tag{S21}$$

$$v_r(\kappa, L) = \frac{1}{\kappa} W_0\left(\frac{V_e^0 \kappa\zeta}{(\zeta+\kappa)} e^{\frac{\kappa(k^- \zeta - E(L-L_0))}{(\zeta+\kappa)}}\right) - \frac{-E(L-L_0) + k^-\zeta}{\zeta+\kappa} \tag{S22}$$

Its phase space is the $L$-$\kappa$-plane, shown in Fig. S6B for a spread model cell ($v_c$=0). The dashed lines in Fig. S6 are called nullclines, and mark those $L$ and $\kappa$ values for which either $dL/dt = 0$ or $d\kappa/dt = 0$ holds. At their intersection point, both dynamics are 0, and the state does not change, i.e., it is an equilibrium state. If the system starts at any state except the equilibrium state, it moves towards the equilibrium state, and draws its trajectory in phase space.

Fig. S6 shows two example trajectories. The green one starts rather close to the stationary state at $\kappa \approx 0.5$ nNsμm$^{-2}$, $L$=20 μm. It quickly moves to the equilibrium state. The orange trajectory starts at $\kappa \approx 1.9$ nNsμm$^{-2}$, $L$=20 μm. It starts on a flux line first leading to much larger $L$ - the protrusion grows. It then turns to very small values of $\kappa$, where it turns parallel to the $\dot{\kappa}$-nullcline, and $L$ decreases even below the stationary value - the protrusion retracts. This retraction is fast, since it happens at small $\kappa$, and corresponds to a slippage event. Slippage events occur whenever the system reaches this low-$\kappa$ branch of the $\dot{\kappa}$ nullcline. Finally, the trajectory approaches the equilibrium state. The large amplitude of the initial protrusion growth and the slippage event set the orange trajectory apart from the green one on a phenomenological level. We introduced a basin of attraction in Fig. 5C, to distinguish the trajectories going directly to the equilibrium state from the trajectories including slippage.

### S9 Force-retrograde flow regimes and direction reversal

Eq. S19 shows that force has a biphasic relation with the retrograde flow in the stationary state. Thus, change of force upon variation of retrograde flow depends on the regime that the protrusion is working in. Retrograde flow is not symmetric in a moving cell. It is the vectorial sum of extension rate and edge velocity (Fig. 5F). This entails higher retrograde flow at the cell back. The right panel in Fig. S7 shows the retrograde flow in the simulations during a direction reversal. In the beginning, the retrograde flow at the back is higher than at the front. This changes after the reversal swapping the roles of front and back. We find that the retrograde flow at the front is always lower than $v_{r,cr}$, and thus working in the rising branch of force - retrograde flow relation (Fig. S7, left). However, on the other side of the cell, back retrograde flow is always higher than $v_{r,cr}$. The faster the cell moves, the larger is the separation between the retrograde flow at the front and back protrusions.

The left panel in Fig. S7 shows that both protrusions in a slow cell work close to the critical retrograde flow rate, $v_{r,cr}$. Reversal of the direction requires switching the roles of front and back. Thus, the reversal should be easier in slow cells, whose protrusions work close to the $v_{r,cr}$. This argument agrees with the lower persistence time and higher reversal probability of slow cells.

Differences in the force-retrograde flow regime lead to a characteristic difference between the front and back protrusions. The retrograde flow rate has positive feedback on force generation at the front. It speeds up at the front, if the front edge motion is slowed down due to back pulling. This increases the force generation at the front, which tends to oppose the velocity reduction caused by the pulling of the back. However, the retrograde flow rate has a negative feedback on the force generation at the back. If the back edge speeds up due to the stronger front pulling, the back retrograde flow also speeds up. As a result, the back force resisting motion decreases. In this sense, the back even helps the front pulling to increase the cell velocity.

To see how the force-retrograde flow regimes change during a reversal, we perturbed the friction coefficient $\kappa$ at the back to induce a cell reversal. Fig. S8A shows a case when the perturbation is not large enough and cannot trigger the reversal. The panel on the bottom shows the evolution of back and front protrusions in the force-retrograde flow plane. The perturbation at the back is not sufficiently large. So, front and back states eventually return to their original locations on the biphasic curve of the steady states. Fig. S8B shows a case with a sufficiently large perturbation that can induce the reversal. The panel on the bottom shows how the back state moves from the falling branch to the rising branch of the biphasic steady state curve. Fig. S8C shows the flux lines in this F-$v_r$ plane. Although they are obtained from the single protrusion model with the assumption that the velocity of the cell body stays unchanged, they can still approximate the behaviour of the full model. The flux lines are shown for a front protrusion (upper panel), a protrusion in the spread state (middle panel), and a back protrusion (bottom panel). In a front protrusion, the flux lines converge to a fixed point on the rising branch of the biphasic steady state force-retrograde flow curve. The fixed point is on the falling branch of the steady state curve in back and spread protrusions.

### S10 Stationary force-velocity relation of cell motion and stall force

To determine the stationary force-velocity relation of cell motion, we include an external force acting on the leading edge membrane of the cell, $F_m$ in our formulation. The force balance at the leading edge gives:

$$F_f - E(L_f - L_0) - \zeta_f v_f - F_m = 0 \tag{S23}$$

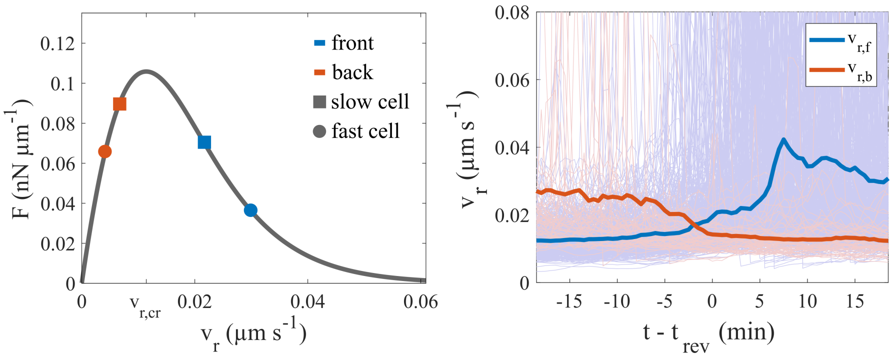


Figure S7: Left: The biphasic relation between force and retrograde flow in the stationary state. $v_{r,cr}$ is the critical flow rate with maximal force. Symbols show the steady state retrograde flow and force at the back and front protrusions for a fast and a slow cell. Right: Retrograde flow at front ($v_{r,f}$) and back ($v_{r,b}$) protrusions during a direction reversal as average over many cell tracks (thin lines). $t_{rev}$ is the time when the cell nucleus changes direction. The results are obtained from simulations similar to Fig. 5A with the parameter values of set 1 listed in Table S2.

With that, Eq. S7 changes to:

$$v_f = \frac{N_f}{a\zeta_f} W_0\left(\frac{V_e^0 a\kappa_f\zeta_f}{N_f(\zeta_f+\kappa_f)} exp\left(\frac{a\kappa_f(k^-\zeta_f - E(L_f-L_0)-F_m)}{N_f(\zeta_f+\kappa_f)}\right)\right) - \frac{E(L_f-L_0)+F_m+k^-\kappa_f}{\zeta_f+\kappa_f}, \tag{S24}$$

We use Eq. S24 instead of Eq. S7 together with all the other equations to find the cell velocity in the stationary state in dependence on $F_m$. Fig. S9 shows velocities and forces in dependence on the external force. We present the deterministic relation without noise here to be comparable to earlier published predictions. Cell velocity decreases with increasing $F_m$, but in two different regimes. At low $F_m$, front retrograde flow speeds up with increasing $F_m$, due to the decrease of the leading edge membrane velocity caused by the opposing external force. At a certain $F_m$ the front retrograde flow comes very close to the critical value $v_{r,cr}$ (Fig. S9, middle). In this situation, the protrusive force at the front $F_f$ reaches the maximum force that the protrusion can produce (see the biphasic force-retrograde flow relation in Fig. S8). $F_f$ cannot increase more after this point by increasing $F_m$ (Fig. S9, right). Thus, the stationary force velocity relation enters a new regime, with saturated protrusive force:

$$v_f = \frac{1}{\zeta_f}\left(\kappa_f v_{r,cr} - (E(L_f-L_0)+F_m)\right) \tag{S25}$$

Increasing $F_m$ in this regime decreases the velocity almost linearly with small deviations from linearity arising from the interaction of all mechanical components. Cell motion stalls when $\kappa_f v_{r,cr} = E(L_f-L_0)+F_m$ is reached.

Mathematical models of cell motion with linear friction of retrograde flow predicted a linear stationary force-velocity relation or piecewise linear relation very similar to Fig. S9 [25, 1]. Here, we find also a monotonously decreasing function (in difference to the non-monotonous dynamic force-velocity relation reported in [26, 27, 18]). Model predictions for the stationary relation agree in the point that this relation reflects the retrograde flow friction

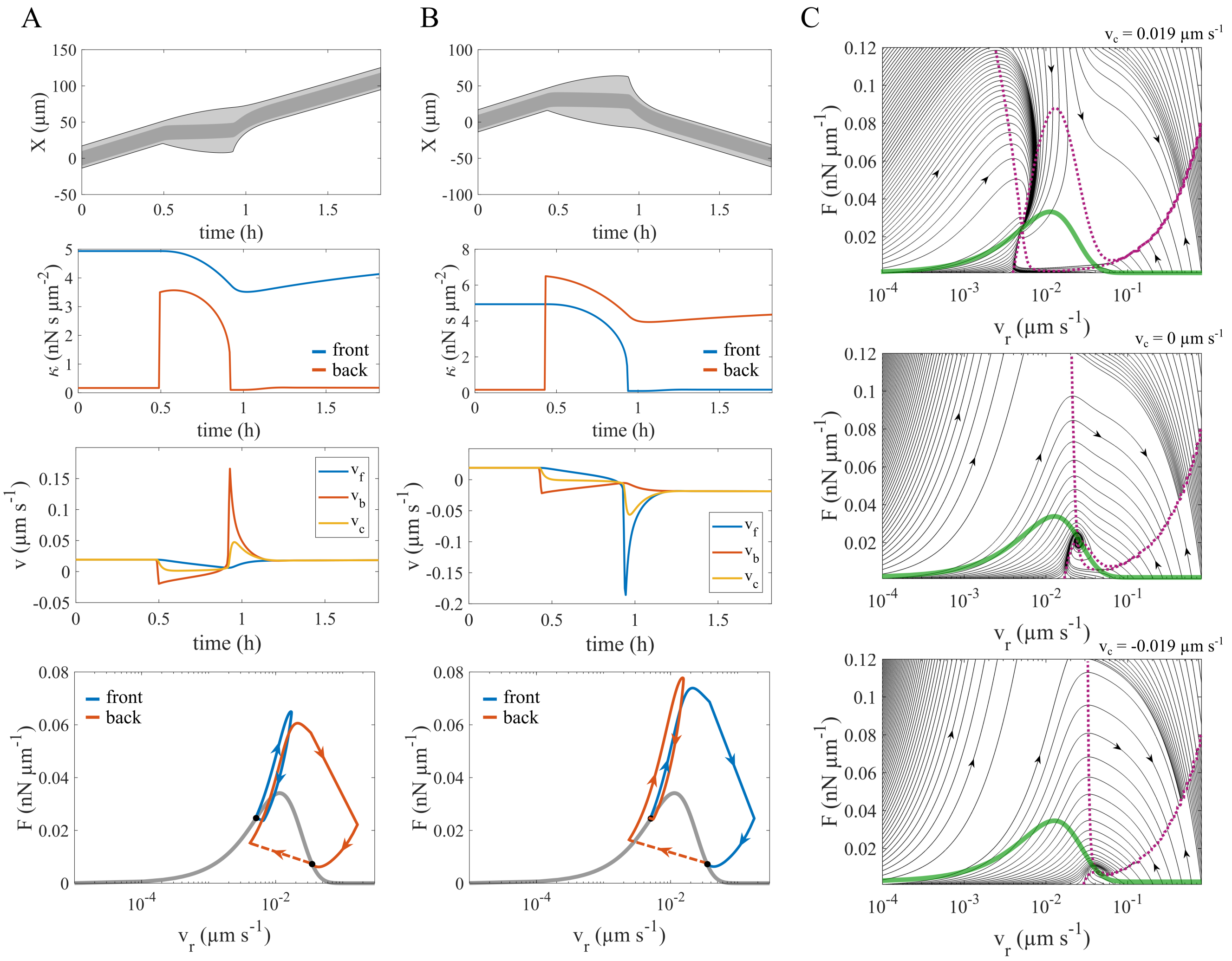


Figure S8: The behaviour of force and retrograde flow at the front and back protrusions during a direction reversal. (A) Cell evolution during a back excitation that does not lead to a reversal. Kymograph, friction coefficient $\kappa$, and velocity of the cell front (blue) and back (red) are shown during the back excitation. The back excitation is started by an instantaneous increase of $\kappa$. The lower panel shows the evolution of front and back in the force-retrograde flow plane. The dashed part of the red line indicates the perturbation of $\kappa$ at the back. The grey line shows the steady state biphasic relation between force and retrograde flow. (B) Cell evolution during a back excitation that leads to a reversal. Kymograph, friction coefficient $\kappa$, velocity, and the trajectory in the force-retrograde flow plane are shown as in (A). (C) Flux lines in the force-retrograde flow plane. They are obtained using the model for a single protrusion (S8) with the positive (up), zero (middle), and negative (bottom) cell velocity. The green line shows the biphasic force-retrograde flow relation in the stationary state. Purple dotted lines show the nullclines of the system in the F-$v_r$ plane. The parameter values of set 1 (Table S2) are used with B = 20 ng cm$^{-2}$ for the simulations in this figure.

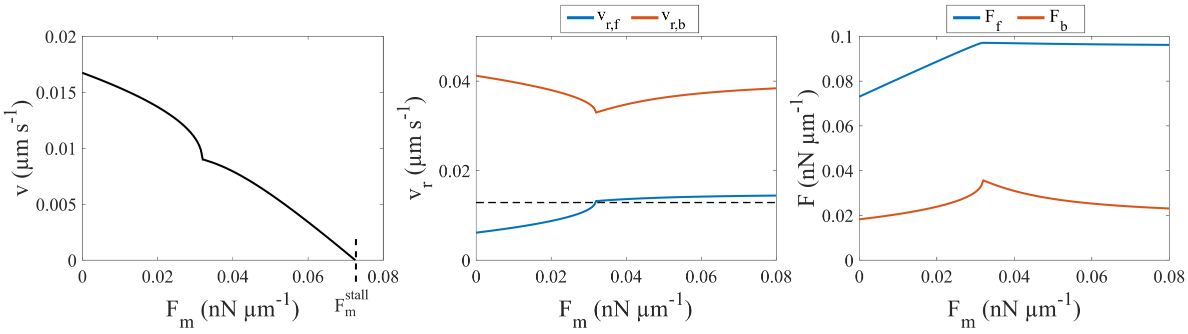


Figure S9: Stationary force-velocity relation. Left: The relation of cell velocity $v$ and external force $F_m$. The cell stalls at the force $\mathrm{F}_\mathrm{m}^\mathrm{stall}$. Middle: Retrograde flow $v_r$ at the front and back protrusions in dependence on $F_m$. The dashed line indicates the critical velocity $v_{r,cr}$ of the relation between friction force and retrograde flow velocity with maximal force (see section S3.) Right: Force at the front and back protrusions $F$ in dependence on $F_m$. The parameter values of set 1 (Table S2) are used in this figure. The Fibronectin density B is assumed to be 45 ng cm$^{-2}$.

law. These model predictions are in agreement with the stalled state of the dynamic force-velocity relation, since the stall force is determined by retrograde flow [18]. The stationary force-velocity relation thus provides access to the 'internal' property retrograde friction law from 'outside'. While the dynamic relation has been measured and analysed in detail [26, 27, 18], the stationary force-velocity relation has not been measured, yet.

### S11 Stimulating direction reversal by Fibronectin steps

In order to substantiate the picture of competing protrusion and the concomitant probabilities of reversals, we designed an experiment where cells experience an abrupt change in the parameter B as a Fibronectin density step. Additional motivation for experiments on heterogeneous lanes arise from the heterogeneous adhesion strength cells moving in organisms during development, metastasis or wound healing experience. MDCK cells may even modify ligand density by Fibronectin deposition and thus spatially constraint their motion [28]. We address these observations by investigating cell behaviour at adhesion steps.

A cell can be perceived as a bistable system within a range of step heights around 0 (Fig. S10A, B). Unlike the homogeneous lanes of Fibronectin, the adhesion density beneath the cell front and back are not equal on the Fibronectin steps. We performed a two-parameter bifurcation analysis on the cell in this situation with parameters $\mathrm{B}_1$ and $\mathrm{B}_2$ being the Fibronectin concentrations forming the step. The Fibronectin density at the cell body $\mathrm{B}_c$ is assumed to be the average of $\mathrm{B}_1$ and $\mathrm{B}_2$.

Fig. S10A shows results of the bifurcation analysis. Cells are in a bistable regime allowing for motion in both directions in the green region. Hence, cells do not necessarily move towards the higher Fibronectin fields, even in the noise-free model. Except on very large Fibronectin steps (purple regions) cells have both up-moving and down-moving stable states. Hence, the concept of multistability also explains the crossing of Fibronectin steps.

The absolute values of the velocity in either direction are not equal with a cell on a step. The speed with which

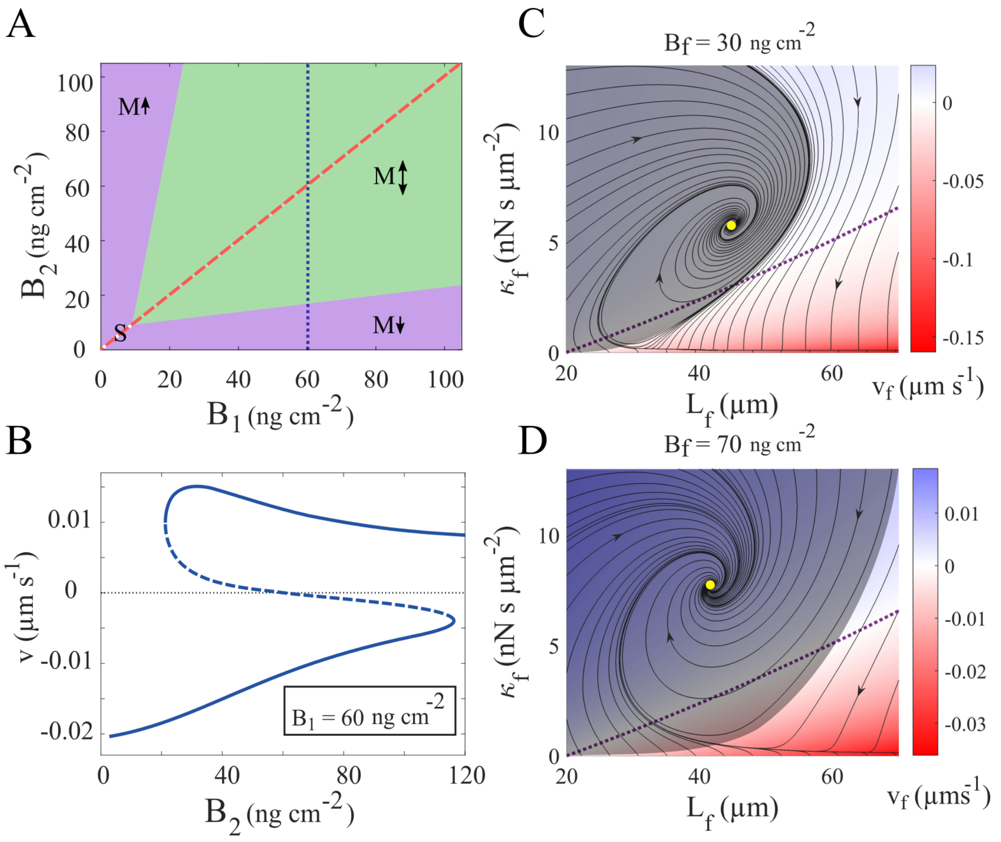


Figure S10: Cell states on FN steps. (A) 2D bifurcation diagram with respect to Fibronectin densities forming the step $B_1$ and $B_2$. Cells can move only in one direction in the purple regions. Both upward and downward states coexist in the green region. (B) Cell states illustrated by their value of the cell velocity for a range of Fibronectin densities at the cell front $B_2$. The Fibronectin density at the cell back $B_1$ is 60 ng cm$^{-2}$. This corresponds to the blue dotted line in panel A. (C), (D) The basin of attraction of the steady protrusion state (grey area) and state trajectories (black lines) in $\kappa$-$L$ plots. The dotted purple lines represent zero velocity. The parameter values of set 4 (Table S2) are used in this figure.

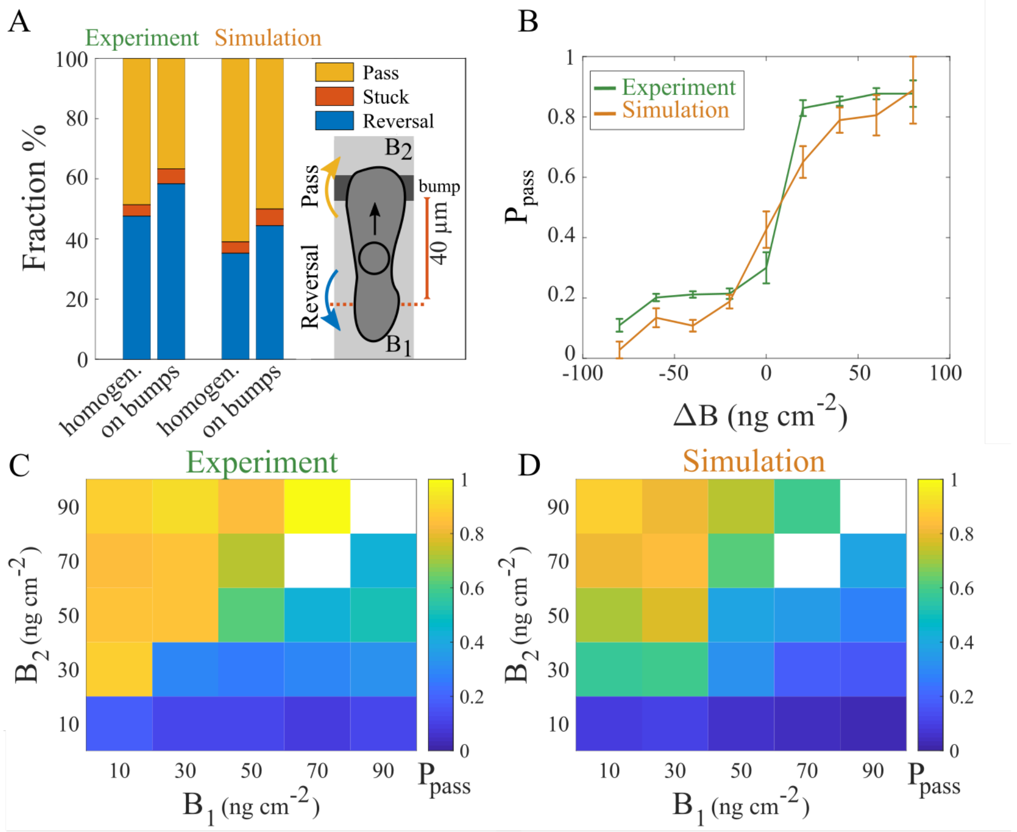


Figure S11: Cell behaviour at Fibronectin steps. (A) Examples for the probabilities of the three choices passing, reversing direction and getting stuck on a homogeneous area and at a Fibronectin 'speed bump' of about 10 ng $cm^{-2}$ and 10 µm width. We observe the nucleus when it enters a 40 µm range before the step. If it leaves the range on the side it entered, it is a direction reversal, if it leaves across the step, it is a pass, and if it remains longer than a dwell time of 5.5 h in this range, it got stuck. The choice of this dwell time is according to our definition of a spread cell. (B) The probability to pass $P_{pass}$ in relation to the step height averaged over a range of initial densities from 0 to 100 ng $cm^{-2}$. (C), (D) $P_{pass}$ for all measured pairs of $B_1$ and $B_2$. Cells move from $B_1$ into $B_2$. White colour indicates data not measured. The parameter values of set 4 (Table S2) are used for the simulations in this figure.

the cell moves into the area with higher adhesion is different from the speed in opposite direction. Applying our concept of the basin of attraction reveals that the front protrusion of a cell moving into a higher-adhesion area is more stable than the one of a cell moving into a lower-adhesion area (Fig. S10C, D).

A cell facing a step has three choices: passing the step, turning around or getting stuck (Fig. S11A). A small Fibronectin step of 10 ng $cm^{-2}$ changes the probabilities for all three choices already. Fig. S11B shows that cells like to move into high-adhesion areas (positive $\Delta$B). This is in agreement with our calculation of larger stability of front protrusions during the transition than during motion in the opposite direction. This preference for high adhesion density is in agreement with haptotaxis and the restriction of cell motion by Fibronectin deposition mentioned above. However, cells are also able to move into low-adhesion areas (negative $\Delta$B), and thus do not get stuck in places with high adhesion.

Remarkably, cells do not pass with certainty even on large steps up the Fibronectin concentration. A small probability for reversal remains. This observation can be comprehended with the concept of the cell on a step being a bistable system where noise may cause transition to either state.

We could again reach good agreement between experimental data and simulations for each of the data sets presented in Fig. S11. Hence, the set of ideas defining the theory is also able to explain the motion of cells across Fibronectin steps.

### S12 Remarks on UCSP simulations

The persistence time is defined as the time the nucleus of a cell moves continuously in the same direction. The *Instantaneous velocity* of the cell has been determined from the nucleus position of consecutive frames without any averaging over several time steps or cutoffs. The *velocity* is the difference of positions of consecutive direction reversions divided by the difference of the points in time of these reversions. The data in Fig. 5B are the average over all the cell trajectories in each condition.

A histogram of the persistence time of the moving episodes shows that the reversion of the direction is a Poisson process. To determine persistence time and the corresponding velocity, we analysed data sets with two different time resolutions (10 min and 30 s). The same temporal resolutions and Fibronectin densities are used in the corresponding simulations to determine persistence time and instantaneous velocity. All other parameters in the simulations are the same as Table S2, set 1. With this definition, persistence time is dependent on the temporal resolution of the cell measurement. The calculated persistence time is shorter for data with higher temporal resolution because occurrence of one small time step with displacement in the opposite direction is more likely than a large time step with reversed displacement. The low temporal resolution of 10 min removes the effect of the small fluctuations of the nucleus position on the determination of persistence length, while the high temporal resolution captures these fluctuations. Different velocities of moving episodes are caused by the noise in the system. We find that there is a positive correlation between cell velocity and the persistence time (Fig. 5B). However, they are exponentially correlated only in the low velocity range. The persistence time and velocity show similar dependency in the drug treatment experiments.

### S13 Ideas of morphodynamic mechanisms in MDA-MB-231 cells discussed in literature

d'Alessandro et al. showed with MDCK epithelial cells that Fibronectin deposited by the cells themselves during motion might generate a range of increased Fibronectin density on the substrate [28]. MDCK cells on 1d Fibronectin lanes like to stay within the region where they increased the Fibronectin density, and only leave it after having turned around several times at its boundaries. d'Alessandro et al. explain their observations by cells shying away from passing Fibronectin steps down. This observation agrees with our results on behaviour of MDA-MB-231 cells on artificial Fibronectin steps. The bistable behaviour of cells on steps explained in section S11 offers a mechanistic explanation. The probability to pass steps towards lower Fibronectin density is small (Fig. S11B), i.e. cells turn around more often than they pass.

Oscillatory motion with a constant or slowly monotonously increasing amplitude, which is larger than the cell size, is the hallmark of the Fibronectin deposition effect on cell trajectories [28]. We did not observe oscillations of this type in our experiments. Oscillations with constant amplitude in our trajectories had amplitudes close to cell size and thus cannot be distinguished from state SO and were classified as such.

Bolado-Carrancio et al. [29] investigate the coordination of the Rho GTPase network activity across MDA-MB-231 cells. The mainly experimental study also simulates a mathematical model and is closely related to Holmes et al. [30]. Holmes et al. [30] favour an oscillation mechanism with bistability resulting from the Rac1-RhoA-antagonism. Integrin signals via paxillin, FAK and Src to activate Rac1 and suppresses RhoA [31, 32]. RhoA and Rac1 are additionally connected in a signalling network also comprising DIA, ROCK and PAK [29]. This network acts in a way that in the end RhoA and Rac1 mutually suppress their activity to a degree controlled by the presence of DIA and/or ROCK. This double negative feedback causes the bistability. Holmes et al. [30] introduced a slow negative feedback from interaction with ECM due to area changes of lamellipodia modulating integrin signalling. The slow feedback turns the bistability into oscillations. Conservation of total Rac1 and RhoA turned out to be essential for robust agreement of RhoA/Rac1 dynamics with experiments [30, 33, 29]. The Rac1-RhoA-antagonism with conservation of total Rac1 and RhoA produces anti-phase oscillations [30, 33], as systems with essential feedback on the basis of resource limitations or conservation have the tendency to do. Park et al. state "Conservation of the total amount of small GTPases in a cell effectively leads to a double negative feedback between the front and the back of the cell, preventing simultaneous high activation in both front and back cellular compartments." [33]. We do not observe strictly antiphase or in-phase oscillations but rather a continuum of phase relations.

Bolado-Carrancio et al. [29] report leading edge Rac1-mediated oscillations with a period in the 30-60 s range at the front protrusion. They suggest the back to be most of the time in a state with high RhoA activity and activated ROCK. That state is interrupted by waves of high Rac1 activity from the front arriving with a period of about 4-5 min at the back (their Figure 5-figure supplement 1C, control). Bolado-Carrancio et al. suggest these waves to cause periodic back retractions.

The cell movement cycle is described by the authors in the introduction as “The leading edge protrudes and retracts multiple times, until the protrusions, known as lamellipodia, are stabilised by adhering to the extracellular matrix (Ridley, 2001). Subsequently, the cell back detaches and contracts allowing the cell body to be pulled toward the front.” The cell movement cycle takes about 45 min (Bolado-Carrancio et al. [29] text and movie 2). Hence, on average one in ten wave arrivals causes retraction of the back.

The retraction mechanism is based on myosin. It is explained as “Because of the oscillations, zones of low Rac1 activities emerge, which give rise to high RhoA-GTP that interacts with ROCK and leads to the back retraction (Video 1). Subsequently, RhoA returns to its initial high stable activity, and the dynamic pattern of RhoA-GTP and Rac1-GTP over the entire cell returns to its initial state. These model simulations could plausibly explain how the different GTPase dynamics at the cell front and back are coordinated to enable successful cell migration.” Here, oscillations at the back are meant and are also called ‘adhesion-retraction cycle at the back’. Details on how the Rac1-activity at the back causes retraction but the state with high ROCK activity activating myosin does not are not provided. The reasoning is, where Rac1 is low, RhoA is high, activates ROCK and this in turn phosphorylates the myosin light chain causing contraction. However, the areas of low Rac1 during the waves are much smaller than during the stationary state and therefore contraction should be weaker. Also and consequently, there should be no retractions with Blebbistatin, if we face this mechanism. Our finding that oscillations and thus protrusion retractions are observed with Blebbistatin applied is difficult to reconcile with the myosin based retraction mechanisms suggested in ref. [29]. We found myosin’s role in formation of adhered structures to be more important than as a contractile driver of F-actin flow and protrusion retraction [1].

We consider the likelihood that the oscillations we observe obey the GTPase-mechanism by looking at time scales. The oscillations that we observe at the back have average periods in the range from 15 min to 30 min, i.e. are slower than the Rac1-wave period and faster than the movement cycle reported by Bolado-Carrancio et al. We do not observe exclusively antiphase oscillations, as the GTPase-mechanism suggests, but a whole range of phase relations. We find similar time scale differences when comparing our observations to the RhoA-RhoGDI-based pacemaker mechanism suggested by Tkachenko et al. [34] acting on a time scale of about 100 s. Interestingly, interrupting the pacemaker function by inhibiting its negative feedback to RhoA by inhibiting PKA increased the typical time scale to about 10 min.
